# Integrated Analysis of Glycosylation and Inflammation-Related Genes for Prognostic Risk Modeling and Immunotherapy Response Prediction in Gastric Cancer

**DOI:** 10.1101/2025.09.15.676447

**Authors:** ZhaZhu Li, Musthaq Ahmed, Tao Xu, Henna Li

**Affiliations:** Department of General Surgery, Gansu Provincial Hospital Affiliated to Wuhan University of Medical College, Xiaogan 432000, Hubei, China; Medical College, Wuhan University of Medical College, Wuhan 430065, Hubei, China; Department of Microbiology, virology and immunology. Fergana Medical Institute of Public Health, Fergana, Uzbekistan; 4 Department of Rehabilitation Medicine, Gansu Provincial Hospital of Traditional Chinese Medicine, Lanzhou, Gansu, China

**Keywords:** Gastric cancer, Inflammation, Glycosylation, Prognostic indicators, Immune infiltration

## Abstract

**Background:** Gastric cancer (GC) continues to be among the most commonly identified cancers worldwide. This study integrates glycosylation and inflammation-related gene features for the first time to construct a prognostic model for gastric cancer, providing new theoretical basis for revealing immune escape mechanisms and personalized treatment strategies.

**Methods:** Transcriptomic and clinical data derived from GC samples were meticulously examined, utilizing resources from The Cancer Genome Atlas (TCGA) and Gene Expression Omnibus (GEO) datasets. Through differential expression analysis, we successfully identified glycosylation and inflammatory-related differentially expressed genes (GANDIRDEGs). To construct a prognostic gene signature, we applied least absolute shrinkage and selection operator (LASSO) analysis in conjunction with Cox regression analysis. Additionally, we performed somatic mutation (SM) along with copy number variation (CNV) analyses, alongside gene ontology (GO) and Kyoto Encyclopedia of Genes and Genomes (KEGG) enrichment analyses. Furthermore, we conducted gene set enrichment analysis (GSEA) along with a comprehensive evaluation of immune infiltration and drug sensitivity.

**Results:** We identified and validated a six-gene (*INHBA, OLR1, ROS1, EPHA5, TACR1, and IL6*) signature, termed GANDIRDEGs, which showed excellent performance in distinguishing overall survival (OS) between high-risk (HR) and low-risk (LR) cohorts. Moreover, we developed a prognostic nomogram utilizing this six-gene signature that provides highly accurate predictions of GC patient outcomes.SM and CNV analyses revealed that *MSR1* had the highest mutation rate among the GANDIRDEGs, with a mutation rate of 5%. GO, KEGG, and GSEA revealed significant associations of each pivotal gene with pathways, including cytokine signaling, the inflammatory response, and apoptosis mediated by CDKN1A through TP53, among various biological functions and signal transduction pathways. Our findings offer a novel gene signature, GANDIRDEGs, that correlated with prognosis, immune infiltration, and therapeutic sensitivity in patients with GC.

**Conclusion:** This study establishes a prognostic signature integrating glycosylation and inflammatory pathways in GC, providing valuable insights into the mechanisms of immune evasion and potential personalized treatment approaches.

## 1 Introduction

In 2022, GC was reported to rank as the fifth most prevalent cancer globally, in terms of both incidence and mortality rates [1]. On the basis of incidence trends and prediction models, it is estimated that 10.0 million new cases and 5.6 million GC-related deaths will occur in China between 2021-2035 [2]. GC patients in China often present with advanced-stage disease and high tumor burden, making early diagnosis and prognosis assessment particularly challenging. Currently, the biomarkers used clinically for GC prognosis prediction are limited, and primarily rely on traditional indicators such as histological grading and TNM staging. Nevertheless, these techniques frequently do not adequately encompass the biological traits of tumors or the requirements for personalized therapy. This study is based on international public databases including TCGA and GEO, covering GC patients from different regions around the world. Developing more sensitive and specific molecular features is of great significance for improving the prognosis assessment and treatment decision-making of GC patients worldwide.

In recent years, research has demonstrated that glycosylation is significantly involved in the processes of cancer development and progression [3]. Glycosylation is a key posttranslational modification involving the attachment of complex sugar chains to proteins [4]. Tumor cells often exhibit abnormal glycosylation patterns, which can influence their proliferation, migration, immune evasion, and other biological behaviors [5]. Specifically, changes in glycosylation in cancer cells are closely associated with alterations in the tumor microenvironment, such as proinflammatory signaling. [6,7]. These findings suggest that glycosylation may also regulate the tumor microenvironment. Additionally, inflammatory responses are closely associated with tumor initiation and progression [8]. Inflammation-associated genes are known to have a substantial effect on the progression of GC, not only by driving inflammatory responses but also potentially by promoting glycosylation changes through effects on the expression of glycosylation-related enzymes, thereby forming an interactive mechanism between inflammation and glycosylation [9]. To our knowledge, no previous studies have integrated glycosylation and inflammation-related gene signatures to construct a prognostic model in gastric cancer. This study innovatively constructed a prognostic risk model based on two types of molecular features through multi omics integration analysis of large samples and multiple cohorts, providing new theoretical support and molecular tools for precise stratification and personalized treatment of gastric cancer.

In summary, the relationship between glycosylation and inflammation-related genes offers novel perspectives on the complex biology of GC and a potential direction for identifying new biomarkers and therapeutic targets, which can provide new insights for the early detection and personalized therapeutic approaches of GC. This study aims to identify glycosylation and inflammation-related differentially expressed genes and construct a prognostic model for gastric cancer using integrative transcriptomic approaches.

## 2 Materials and methods

### 2.1 Data acquisition and preprocessing

We acquired stomach adenocarcinoma (STAD) data from The Cancer Genome Atlas (TCGA) (https://portal.gdc.cancer.gov/) via the R package TCGAbiolinks (version2.36.0) [10]. Following the removal of samples that lacked accompanying clinical data, we compiled a total of 352 TCGA STAD samples alongside 30 control samples (normal adjacent tissue), presented in count format. These data were subsequently normalized to Fragments Per Kilobase per Million (FPKM) format. The corresponding clinical data were retrieved from the UCSC Xena database[11] (https://xena.ucsc.edu/). Comprehensive details are provided in Table 1.

**Table 1.** GEO microarray chip information.

|  | TCGA-STAD | GSE19826 | GSE13861 |
| --- | --- | --- | --- |
| Platform | / | GPL570 | GPL6884 |
| Species | / | Homo sapiens | Homo sapiens |
| Tissue | Gastric Cancer Tissue | Gastric Cancer Tissue | Gastric Cancer Tissue |
| Samples in STAD/GC group | 352 | 12 | 65 |
| Samples in Normal/Control group | 30 | 12 | 19 |
| Reference | / | PMID: 21132402 | PMID: 21447720<br>PMID: 29725014<br>PMID: 23112551 |

GEO, Gene Expression Omnibus; GC, gastric cancer; STAD, stomach adenocarcinoma.

Using the R package GEOquery (version 2.76.0) [12], we also downloaded the GC dataset from the GEO dataset [13] (https://www.ncbi.nlm.nih.gov/geo/) GSE19826[14] and GSE13861[15–18]. The chip platform for the GSE19826 dataset was GPL570, which included 12 GC samples and 12 normal samples. The chip platform for the GSE13861 dataset was GPL6884, which included 65 GC samples and 19 normal samples. The detailed clinicopathological parameters are provided in Table 1.

Using the keyword “Inflammatory”, we searched the MSigDB database and identified a gene set containing 200 inflammation-related differentially expressed genes (IRGs). Additionally, we retrieved 282 glycosylation-related genes (GRGs) from published literature in PubMed(https://pubmed.ncbi.nlm.nih.gov/). After removing duplicates, a total of 410 glycosylation and inflammation-related differentially expressed genes (GANDIRGs) were obtained. Detailed information is provided in Table S1.

ComBat method from the R package sva(version 3.56.0)[19] was used to eliminate batch effects from the datasets between merged datasets, yielding the integration of GEO datasets. The merged datasets included 77 GC samples and 31 control samples. The R package limma3.64.0[20] was subsequently used to normalize the integrated GEO datasets, annotate probes, and normalize the data. Principal component analysis (PCA) [21] was performed to assess batch effects between merged datasets.

### 2.2 Identification of differentially expressed genes related to glycosylation and inflammation

The samples were classified into STAD and control categories according to the TCGA-STAD sample classification. The identification of differentially expressed genes (DEGs) was conducted with the criteria of |log2FC|>1, and a false discovery rate (FDR)<0.05. Specifically, genes whose logFC > 1 and adj.p < 0.05 were classified as significantly upregulated. Conversely, genes whose logFC < -1 and adj.p < 0.05 were categorized as significantly downregulated genes. The p-value correction method used was Benjamini-Hochberg (BH). The results of the differential analysis were represented through volcano plots generated with the R package ggplot2 (version 3.5.2.) This was accomplished by intersecting the differentially expressed genes (DEGs) from the TCGA-STAD dataset (with criteria of |logFC| > 1 and adjusted P-value < 0.05) with GANDIRGs, followed by the construction of a Venn diagram. A heatmap illustrating the top 20 GANDIRDEGs was generated via the R package pheatmap (version1.0.12).

### 2.3 Somatic mutation (SM) / copy number variation (CNV) analysis

To investigate SM in the TCGA-STAD cohort, “Masked Somatic Mutation” data were selected as the SM data for the GC cohort in the TCGA-STAD cohort. The data were preprocessed via VarScan (version 2.4.6) software, while visualization of the SM data was achieved with the R package maftools (version2.24.0) [22]. Moreover, to investigate copy number variations (CNVs) within the TCGA-STAD cohort, the “Masked Copy Number Segment” dataset was chosen as the CNV data representative of the gastric cancer (GC) group in TCGA-STAD. Following this selection, an analysis was conducted via GISTIC2.0 [23] on the processed CNV segments, utilizing the default settings.

### 2.4 Gene Ontology (GO) and pathway analysis

Gene Ontology (GO) analysis [24] represents a widely adopted methodology for extensive functional enrichment investigations, and includes three primary categories: cellular component (CC), biological process (BP), and molecular function (MF). The Kyoto Encyclopedia of Genes and Genomes (KEGG) [25] is a widely utilized database that provides comprehensive data concerning genetic sequences, metabolic pathways, medical conditions, and pharmacological agents. The GO and KEGG enrichment analyses for the GANDIRGs were conducted via the R package clusterProfiler (version4.16.0) [26]. The parameters for screening included adj.p < 0.05 and FDR < 0.05, and the Benjamini-Hochberg (BH) method was applied for the correction of p-values.

### 2.5 Gene Set Enrichment Analysis (GSEA) [27]

Genes from TCGA-STAD were first ranked on the basis of their logFC values between the GC group and the control group. GSEA was then performed on the entire set of genes in the TCGA-STAD cohort via the R package clusterProfiler (version4.16.0). The analysis was conducted utilizing a random seed of 2020, establishing a minimum gene set size of 10 and a maximum of 500. For GSEA, c2 gene sets were obtained from the Molecular Signatures Database (MSigDB). The criteria for screening were determined on the basis of adjusted p-values and FDR< 0.05, employing the Benjamini-Hochberg (BH) procedure for the correction of p-values.

### 2.6 Development of a prognostic risk model and survival analysis of patients with GC

We conducted univariate Cox regression analysis on GANDIRDEGs with the R package ‘survival’ to develop a prognostic risk model for GC. Only variables with a p < 0.10 from the univariate Cox regression analysis were selected for the subsequent analysis. The outcomes of the analyses were represented visually through the use of a forest plot.

Subsequently, LASSO regression analysis was performed on the GANDIRDEGs that were included in the univariate Cox regression analysis via the R package glmnet (version4.1-8) [28], with the parameter family set to “cox”. This approach effectively addresses the issue of overfitting by integrating a penalty term (lambda multiplied by the absolute value of the coefficients), which subsequently enhances the model’s capacity for generalization. The results of the LASSO regression analysis were visualized via a prognostic risk model diagram and a variable trajectory diagram.

We then conducted a multivariate Cox regression analysis on the GANDIRDEGs identified through LASSO regression to pinpoint genes associated with the prognostic risk model, and we visualized these results using a nomogram.

GC samples from the TCGA dataset were stratified into HR and LR groups according to the median LASSO risk score. The LASSO risk score was calculated via the following formula:

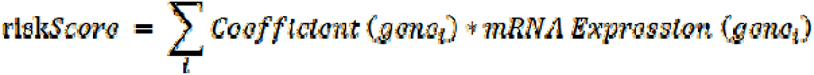

To assess the diagnostic performance of the LASSO risk score in predicting overall survival (OS), a Kaplan–Meier (KM) curve[29] analysis was conducted via the R package “survival”, and KM curves were plotted on the basis of the LASSO risk score. The time-dependent receiver operating characteristic (ROC) curve [30] serves as an important graphical instrument for assessing the efficacy of models, optimizing threshold settings, and comparing different models. Time-dependent ROC curves were constructed via the R package timeROC (version0.4) to calculate the area under the curve (AUC) on the basis of the LASSO risk score and OS, predicting 1-, 2-, and 3-year survival outcomes.

On the basis of the HR and LR grouping of the GC samples in the dataset, the R package ggplot2 was utilized to generate a comparative map to illustrate the different expression levels of genes associated with the prognostic risk model across the two groups. Finally, the risk score for the GC samples in the integrated GEO datasets was derived on the basis of the LASSO risk coefficient. Utilizing the median risk score, the GC samples were categorized into HR and LR cohorts. The same method was used to draw a comparative map showing differences in the expression of the model genes between the two groups in the integrated GEO datasets.

### 2.7 Validation of the prognostic risk model for GC

We performed univariate and multivariate Cox regression analyses to investigate the relationships between the LASSO risk score and clinical outcomes. The results were visualized via a forest plot, which shows the risk score and clinical variables, including age, sex, and clinical stage. A nomogram[31] was generated to represent the functional connections between various independent variables in the multivariate Cox regression model. A calibration curve was constructed to evaluate the precision and differentiation of the prognostic risk model derived from the nomogram. The clinical applicability of the prognostic risk model for survival outcomes at 1-year, 2-year, and 3-year intervals was assessed through decision curve analysis (DCA), utilizing the R package ggDCA (version1.1).

### 2.8 Analysis of immune infiltration

Single-sample gene set enrichment analysis(ssGSEA)[32]was employed to measure the relative presence of various immune cell infiltrates. Each infiltrating immune cell type, including activated CD8+ T cells, regulatory T cells (Tregs), natural killer cells, gamma-delta T cells, and activated dendritic cells, was labeled. The enrichment scores derived from the ssGSEA were used to quantify the relative abundance of each immune cell infiltrate in each sample, creating an immune cell infiltration matrix for TCGA-STAD. The R package ggplot2 was utilized to create comparative maps that illustrate the variations in immune cell expression across distinct groups within the TCGA-STAD dataset. We selected immune cells whose characteristics were notably different between the experimental and control groups for further analysis. Using the Spearman algorithm, we assessed the relationships among immune cells and generated a correlation heatmap to visualize the results. The relationships between model genes and immune cell populations were assessed via the Spearman correlation coefficient. A correlation bubble plot was subsequently produced via the R package ggplot2 to visually represent the findings of the correlation analysis between the model genes and immune cells.

### 2.9 TIDE, MSI, TMB, immune checkpoint and immunophenoscore (IPS) analysis

To evaluate the sensitivity of GC samples to immunotherapy, the tumor immune dysfunction and exclusion (TIDE) algorithm [33, 34] was employed to analyze the acquired dataset. The TIDE immunoscore serves as a potential predictor of tumor treatment response and can be used to assesses the impact of gene expression levels on the overall survival of patients. The differences in the TIDE immunoscore, microsatellite instability (MSI), and tumor mutational burden (TMB) scores between the HR and LR cohorts were analyzed via the Mann-Whitney U test.

Immune checkpoint genes (ICGs), which consist of ligand-receptor pairs, play critical roles in either inhibiting or promoting immune responses. Therapies that inhibit immune checkpoints have demonstrated considerable therrapeutic advantages in various solid tumors. Therefore, we evaluated the expression disparities of 47 ICGs between the HR and LR groups and constructed comparative plots for these groups.

The Cancer Immunome Atlas (TCIA) database[35] (https://tcia.at/home) provides immunophenoscore (IPS) data across 20 different cancer types, and serves as a robust predictor of responsiveness to CTLA-4 and PD-1 therapies. The IPS data for GC samples were obtained from the TCIA database. Comparative plots illustrating the IPS between HR and LR cohorts were subsequently created via the R package ggplot2.

The calculation of IPS score is based on specific gene expression patterns that are closely related to immune cells and their functions in the tumor microenvironment, typically including genes related to processes such as tumor antigen presentation, effector T cell activation, immune checkpoint regulation, and tumor immune escape. In our analysis, the value range of IPS is 0 to 10, with higher scores indicating stronger immunogenicity of the tumor, displaying richer immune cell infiltration and stronger immune response ability.

### 2.10 Drug sensitivity analysis

The Genomics of Drug Sensitivity in Cancer (GDSC) database[36] (https://www.cancerrxgene.org/) is a key public resource for assessing cancer drug sensitivity and identifying molecular markers associated with drug response. The Cancer Cells Line Encyclopedia (CCLE) database[37] (https://sites.broadinstitute.org/ccle/) contains gene expression, chromosome copy number, and large-scale parallel sequencing data from 947 human cancer cell lines, including pharmacological features of 24 anticancer drugs spanning 479 strains, allowing the identification of drug sensitivity predictors on the basis of genetics, lineage, and gene expression. The CellMiner database[38, 39] (https://discover.nci.nih.gov/cellminer/) provides experimental data and gene-drug interaction information for in-depth analysis. We performed an analysis of drug sensitivity concerning pivotal genes, utilizing the expression levels of model-associated genes alongside pharmacological data sourced from the GDSC, CCLE, and CellMiner databases, and the results are presented. The Spearman correlation analysis statistical method was used, and the significance test for all gene drug pairs was conducted using the Benjamini Hochberg method for multiple corrections, with a screening criterion of FDR<0.05.

### 2.11 Statistical analyses

All data processing and analyses were conducted via R software (version 4.2.2). To facilitate the comparison of continuous variables across two separate groups, statistical significance was determined through the independent Student’s t-test for normally distributed variables. In contrast, the Mann-Whitney U test was applied to nonnormally distributed variables. The Kruskal-Wallis test was employed for comparisons involving more than two groups. Spearman correlation analysis was employed to calculate correlation coefficients among various molecules. The statistical p-values were two-sided unless otherwise noted, with a p-value threshold of < 0.05 considered statistically significant.

## 3 Results

### 3.1 Technology Roadmap

### 3.2 Merging of gastric cancer datasets

We used the ComBat method from R package sva to eliminate batch effects from the

GSE19826 and GSE13861 datasets, leading to combined GEO datasets. The distribution boxplot (Fig. 2A-B) and principal component analysis (PCA) plot (Fig. 2C-D) were used to compare the different expression values and low-dimensional feature distributions before and after the elimination of batch effects. The results indicated that the batch effects were effectively removed.

**Fig. 1.**
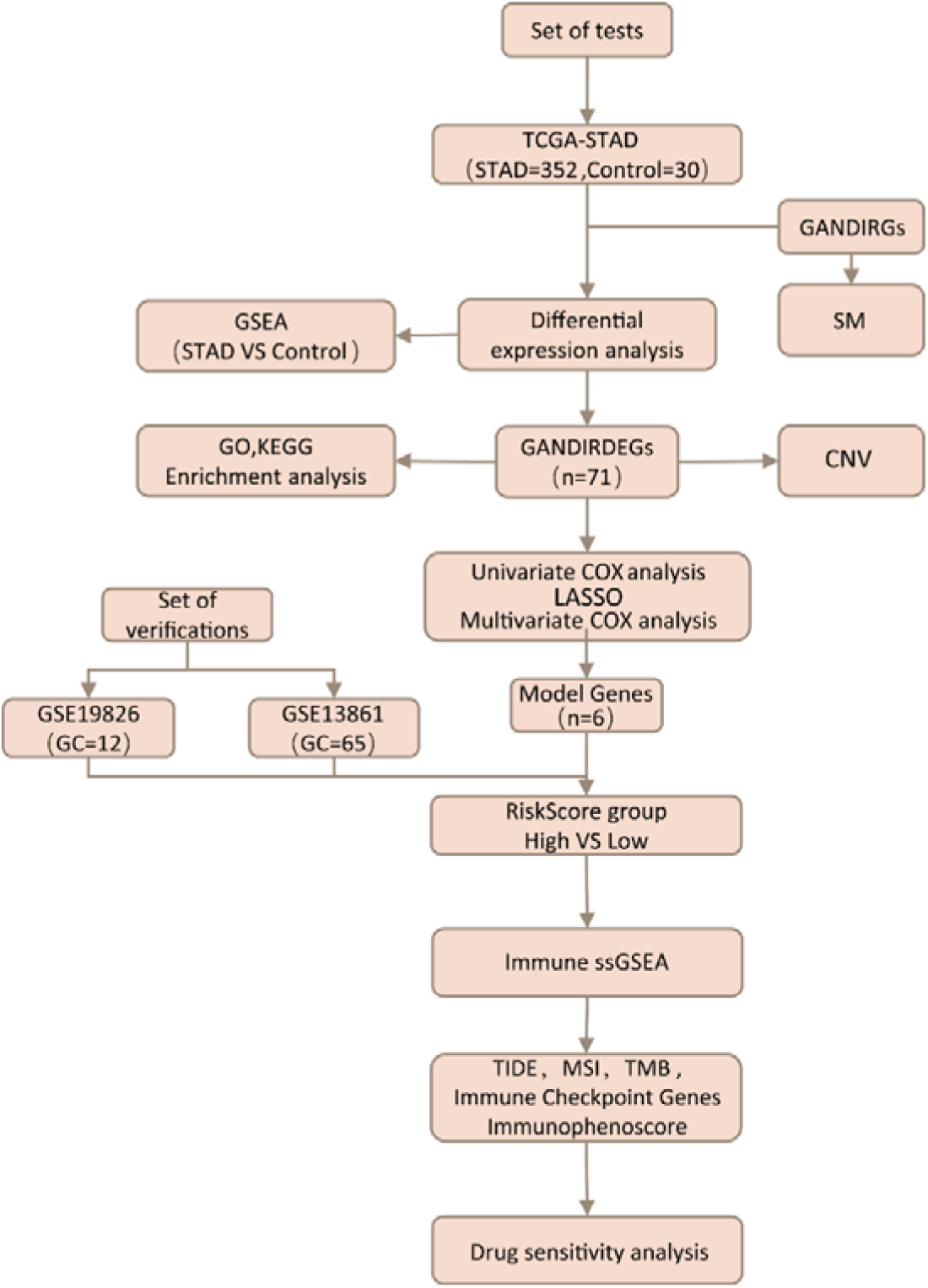
Analytical workflow

**Fig.2.**
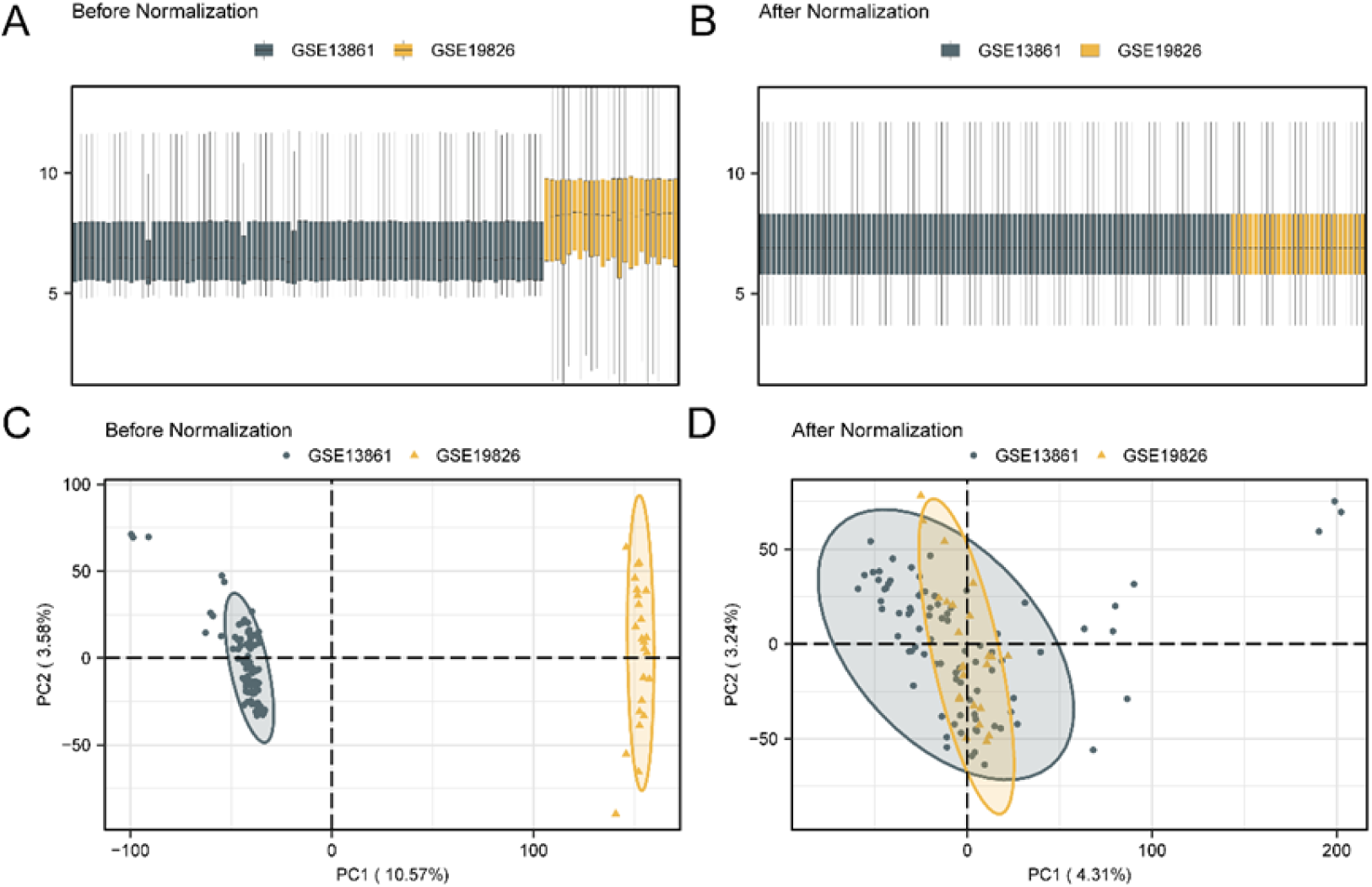
Batch effects removal of GSE19826 and GSE13861. **A.** Distribution boxplot of combined GEO datasets before batch processing. **B.** Combined GEO dataset distribution boxplots after removing the batch effect. **C.** PCA plot of the datasets before removing the batch effect. **D.** PCA plot of combined GEO datasets after batch processing. The GC dataset GSE19826 is in dark yellow, and the GC dataset GSE13861 is in dark green.

### 3.3 Identification of Gastric Cancer-Related Glycosylation and Inflammation-Related Differentially Expressed Genes

The data from the TCGA-STAD dataset were stratified into GC and control cohorts. Differential expression analysis was conducted via the R package limma, which identified 10,070 differentially expressed genes (DEGs) with |logFC| > 1 and adj. P < 0.05. Notably, 6,578 of these genes were determined to be upregulated, whereas 3,492 genes were downregulated. The results were visualized via a volcano plot (Fig. 3A).

**Fig.3.**
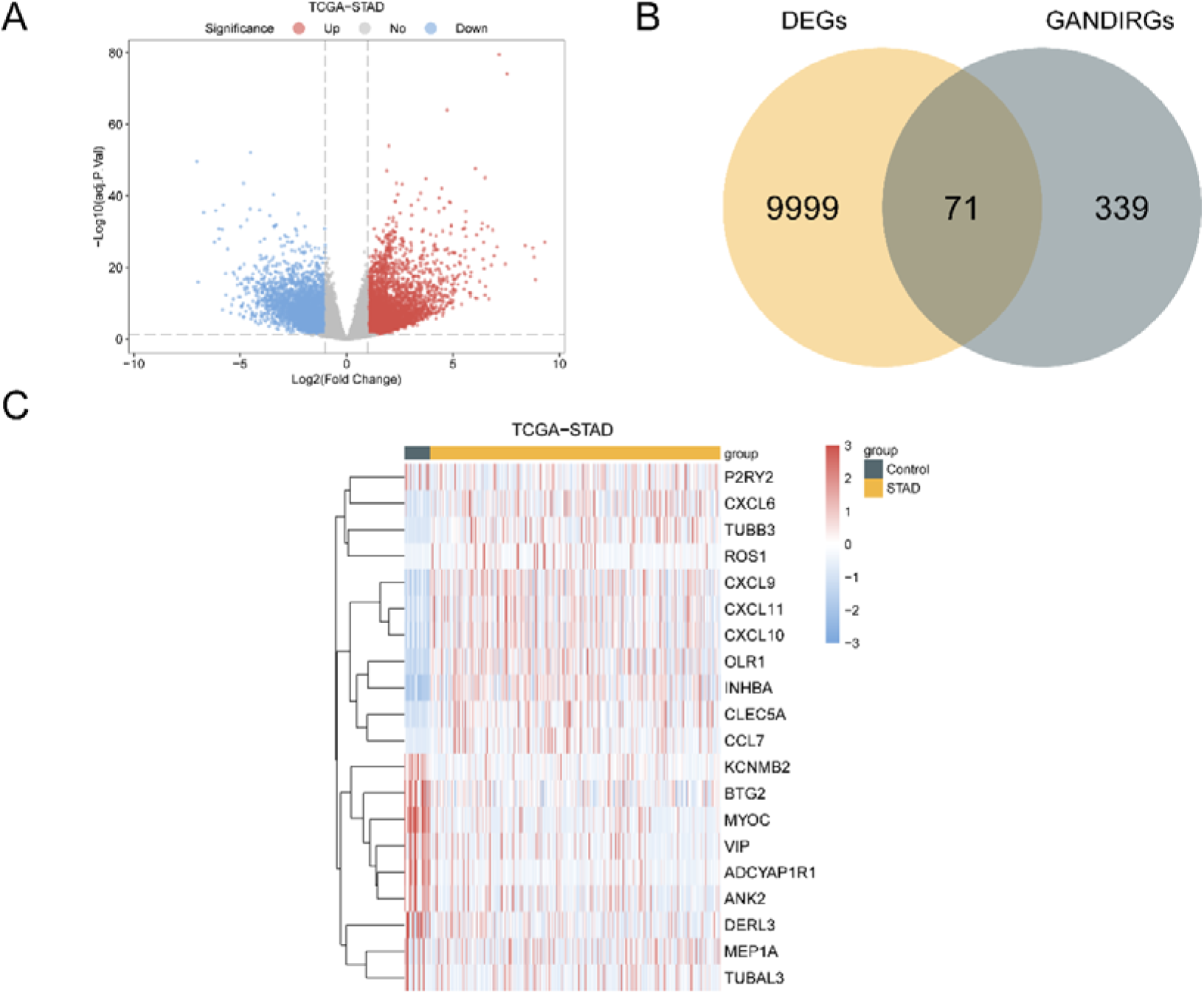
Analysis of Differential Gene Expression**. A.** Volcano plot of differentially expressed genes between the STAD group and the control group in the TCGA-STAD cohort. **B.** Venn diagram of DEGs and GANDIRGs in TCGA-STAD. **C.** Heatmap of GANDIRDEGs in TCGA-STAD. Dark yellow represents the STAD group, and dark green represents the control group. In the heatmap, the color red indicates elevated expression levels, whereas the color blue signifies diminished expression levels.

To obtain GANDIRDEGs, we intersected the DEGs with GANDIRGs and constructed a Venn diagram (Fig. 3B). A total of 71 GANDIRDEGs were identified (Table S2). The top 20 GANDIRDEGs were visualized using a heatmap (Fig. 3C).

The glycosylation and inflammation-related DEGs identified in this study provide a theoretical basis for further exploring the molecular mechanisms and potential therapeutic targets of gastric cancer occurrence and development.

### 3.4 Analysis of gene copy number variation (CNV) and somatic mutations (SM) related to glycosylation and inflammation

To analyze the CNVs of 71 GANDIRDEGs in the TCGA-STAD group, we downloaded and combined the CNV data. GISTIC2.0 analysis identified CNVs in 68 GANDIRDEGs, and the mutation status of the 20 genes with the highest frequency of CNVs was visualized (Fig. 4C-D) (Table S3).

**Fig.4.**
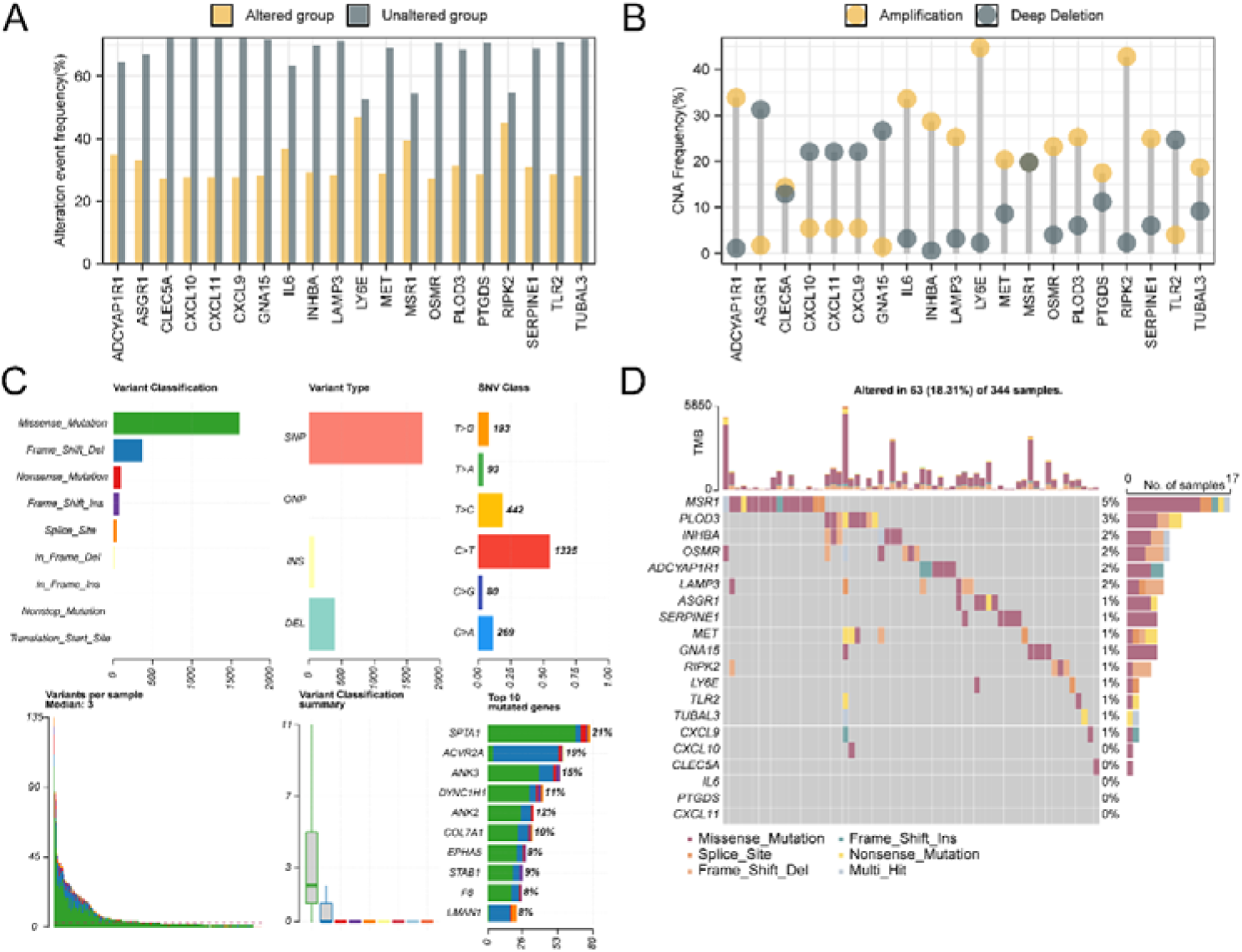
CNV and SM analysis. **A.** SM of GANDIRDEGs in TCGA-STAD. **B.** SMs of the top 20 GANDIRDEGs with the highest mutation frequency in CNVs are shown in the GC group of TCGA-STAD patients. **C-D.** The top 20 GANDIRDEGs with the highest mutation frequency are shown for CNVs in the GC group of the TCGA-STAD cohort.

To analyze the SM among 410 GANDIRDEGs in the TCGA-STAD group, mutation analysis results were compiled and visualized via the R package maftools (Fig. 4A). The analysis indicated the prevalence of nine principal types of SMs among the GANDIRDEGs, with missense mutations being the most frequently observed. Furthermore, the main mutations types of 410 GANDIRDEGs was single nucleotide polymorphisms (SNPs), with C to T mutations being the most prevalent variation observed among single nucleotide variants (SNVs). Visualization of the top 20 GANDIRDEGs with the highest mutation frequency in CNVs (Fig. 4B) revealed that MSR1 had the highest mutation rate among the GANDIRDEGs, with a mutation rate of 5%.

### 3.5 Gene Ontology (GO) and Kyoto Encyclopedia of Genes and Genomes (KEGG) enrichment analysis

To better understand the biological roles of these genes, we performed GO and KEGG pathway enrichment. GO and KEGG enrichment analyses were performed to elucidate the biological processes (BP), cellular components (CC), molecular functions (MF), and biological pathways associated with the 71 GANDIRDEGs in the TCGA-STAD cohort. The comprehensive results can be found in Table 2. The analyses demonstrated that the 71 GANDIRDEGs were enriched primarily in processes related to myeloid leukocyte migration, leukocyte migration, cytokine-mediated signaling pathways, and chemokine-mediated signaling pathways, as well as positive regulation of leukocyte migration and other biological processes. In terms of cellular components, enrichment was observed in the rough endoplasmic reticulum, the external side of the plasma membrane, coated vesicles, membrane rafts, and membrane microdomains, among others. The results also indicated significant involvement in cytokine activity and related molecular functions.

**Table 2.**
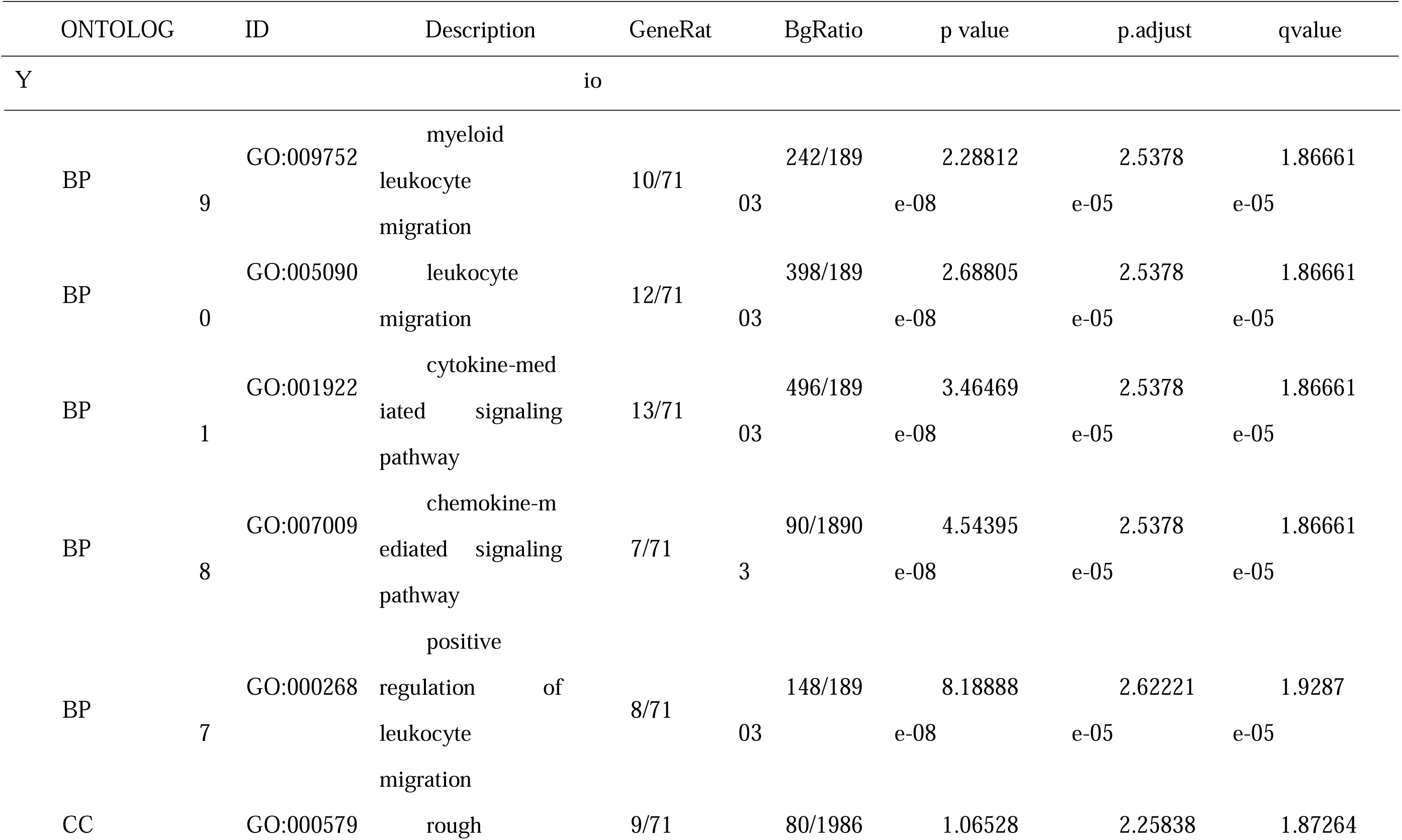
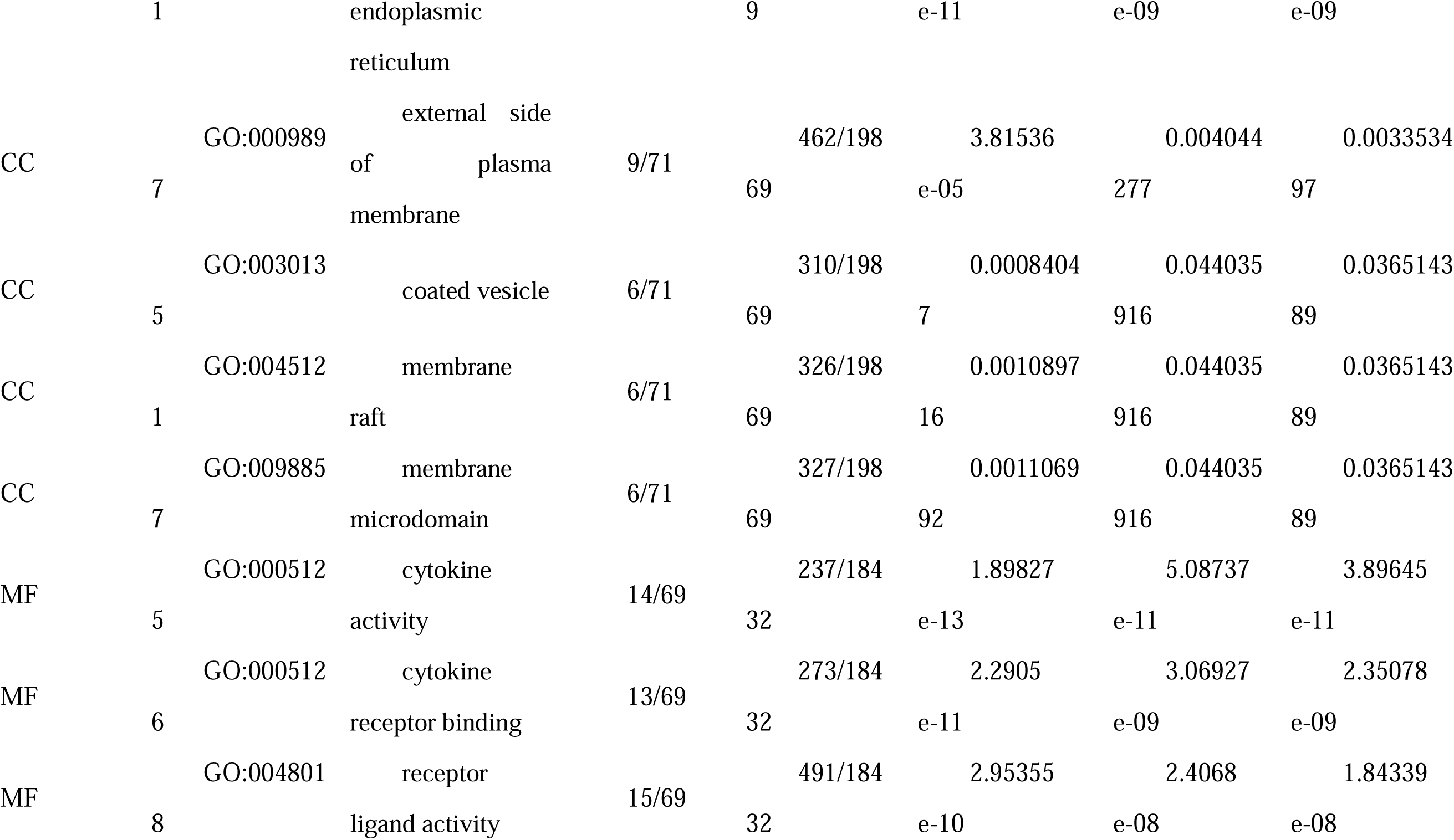
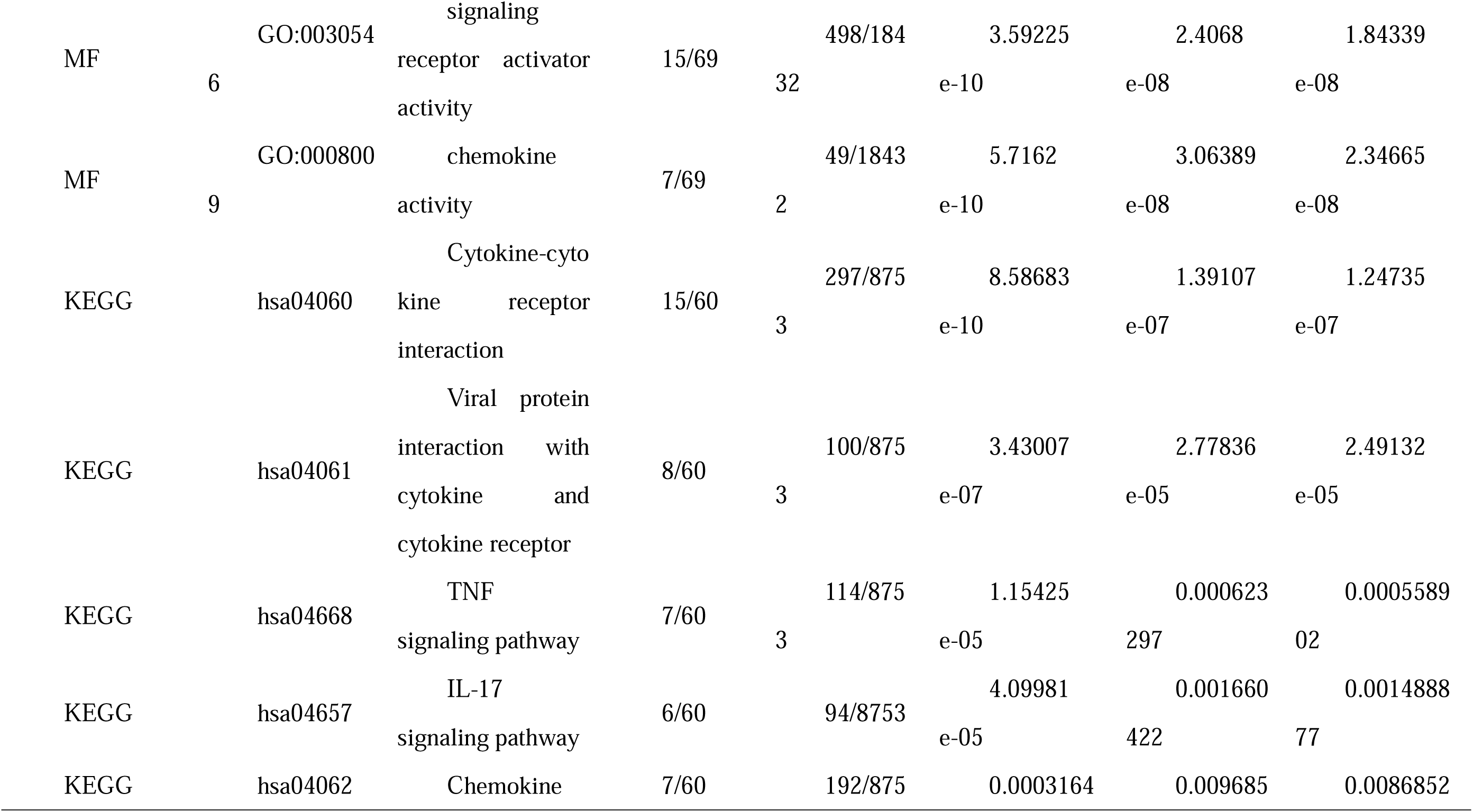
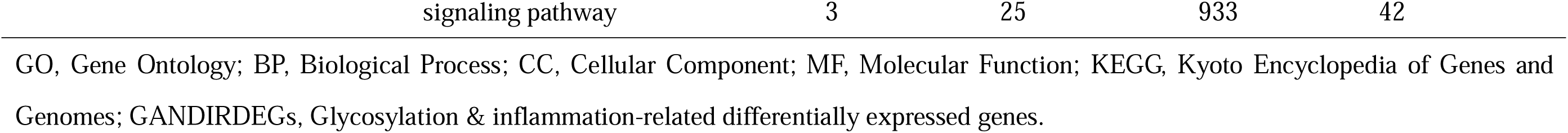
Results of the GO and KEGG enrichment analysis for GANDIRDEGs.

This study identified several molecular functions (MFs), such as cytokine receptor binding, receptor ligand activity, signaling receptor activator activity, and chemokine activity. Furthermore, the investigation revealed substantial enrichment of diverse biological pathways, such as interactions between cytokines and their receptors, as well as interactions of viral proteins with cytokines and their respective receptors, the TNF signaling pathway, the IL-17 signaling pathway, and the chemokine signaling pathway. The results obtained from the GO and KEGG enrichment analyses are presented in bar charts. (Fig. 5A). Moreover, network diagrams for the BP, CC, MF, and KEGG pathways were constructed through enrichment analyses (Fig. 5B-E). In these diagrams, lines represent the respective molecules and their annotations, whereas larger nodes correspond to a greater number of molecules associated with those entries.

**Fig.5.**
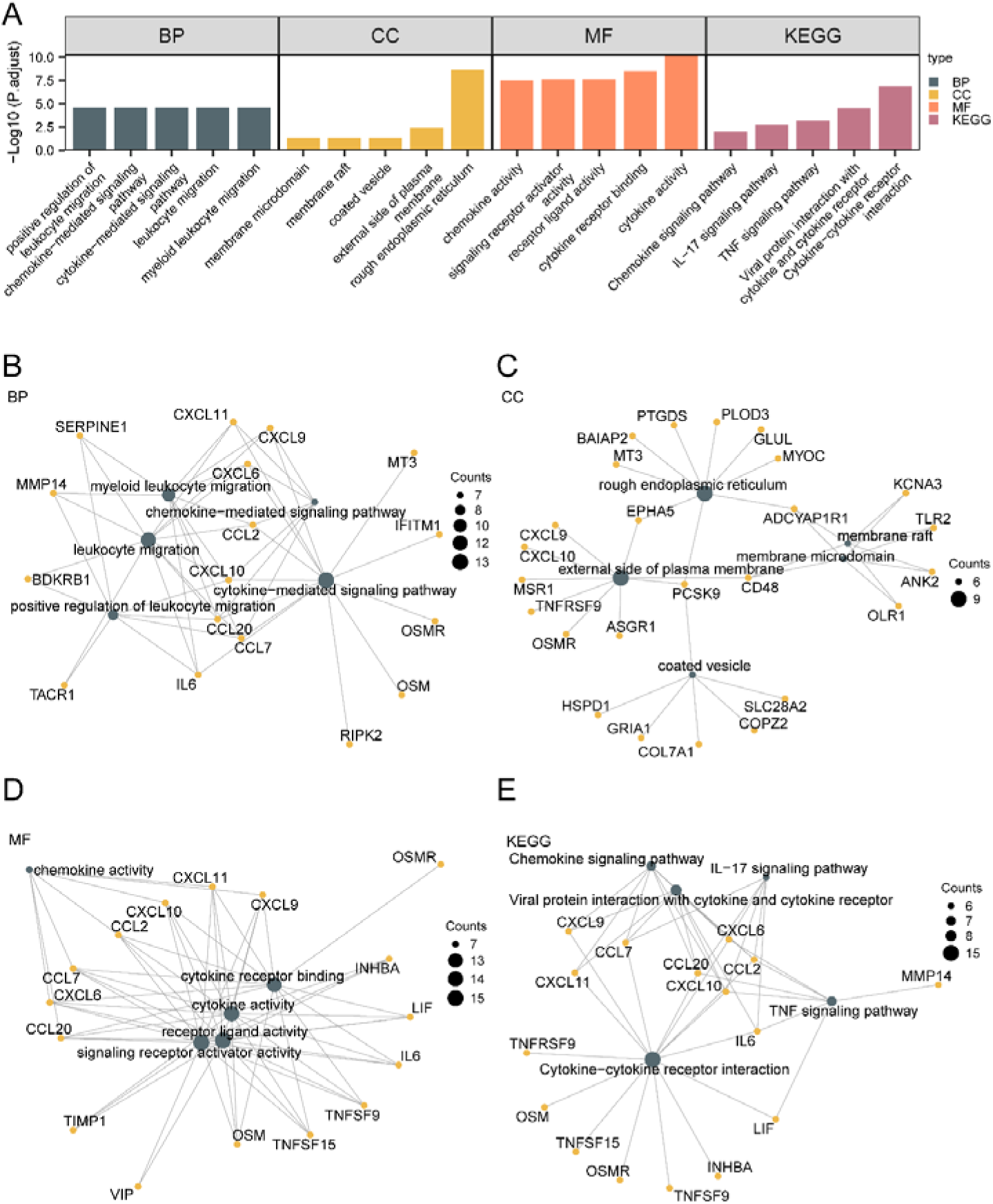
GO and KEGG enrichment analysis for GANDIRDEGs. **A.** Bar graph of gene ontology (GO) and pathway (KEGG) enrichment analysis results of GANDIRGs: biological process (BP), cellular component (CC), molecular function (MF) and biological pathway. The GO and KEGG terms are shown on the abscissa. **B-E.** The results of the GO and KEGG enrichment analyses are shown in the network diagram: BP (B), CC (C), MF (D) and KEGG (E). Dark green nodes denote items, yellow nodes denote molecules, and the lines denote the relationships between items and molecules. The screening criteria for GO and KEGG enrichment analysis were adj.p < 0.05 and FDR value (q value) < 0.05, and the p value correction method was Benjamini-Hochberg (BH).

### 3.6 Gene Set Enrichment Analysis (GSEA)

To assess the impact of gene expression levels on the incidence of STAD, GSEA was conducted utilizing the logFC values of all genes from TCGA-STAD, comparing the STAD group with the control group. This investigation explored the relationships between gene expression levels and a range of biological processes, affected cellular components, and molecular functions, with the results depicted as enrichment plots (Fig. 6A), and summarized in Table 3. The GSEA results indicated that all genes in TCGA-STAD were enriched in various biologically related functions and signaling pathways, including croonquist IL6 deprivation (Fig. 6B), apoptosis via cdkn1a through tp53 (Fig. 6C), the nakamura cancer microenvironment (Fig. 6D), cytokine signaling, and the inflammatory responses (Fig. 6E).

**Fig. 6.**
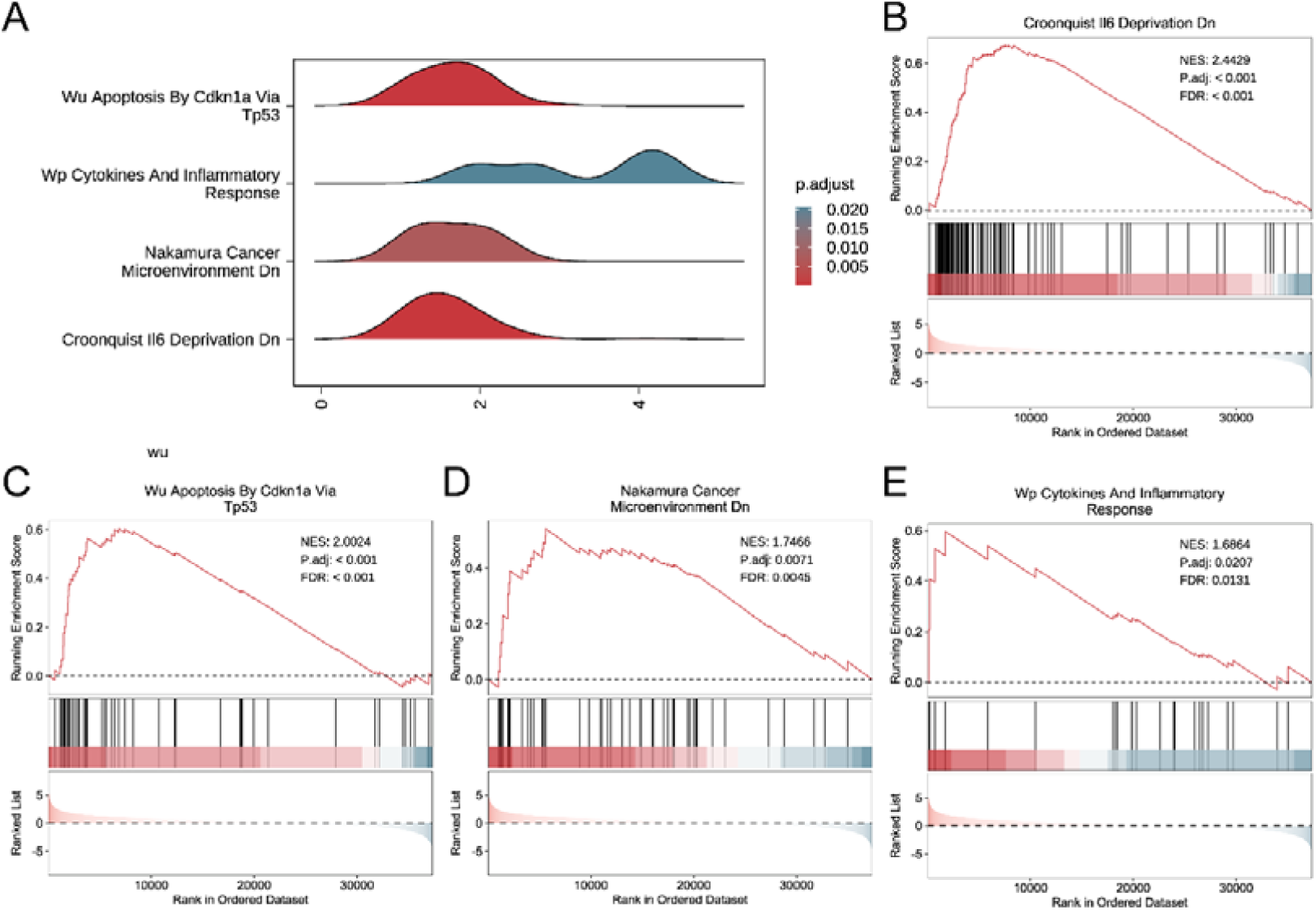
Differential gene expression analysis and GSEA for TCGA-STAD. **A.** Mountain map displaying the 4 biological functions of GSEA in TCGA-STAD. **B-E.** GSEA showed that the TCGA-STAD cohort was significantly enriched in croonquist IL6 deprivation (B), wu apoptosis by cdkn1a via tp53, the nakamura cancer microenvironment dn, cytokines and the inflammatory response (E). In the enrichment plot, the color represents the size of the adj.p value, the redder the adj.p value is smaller, and the bluer the adj.p value is larger. The criteria employed for screening in GSEA included adj.p < 0.05 and FDR value (q value) < 0.05. The method utilized for p-value correction was the Benjamini-Hochberg (BH) procedure.

**Table 3.**
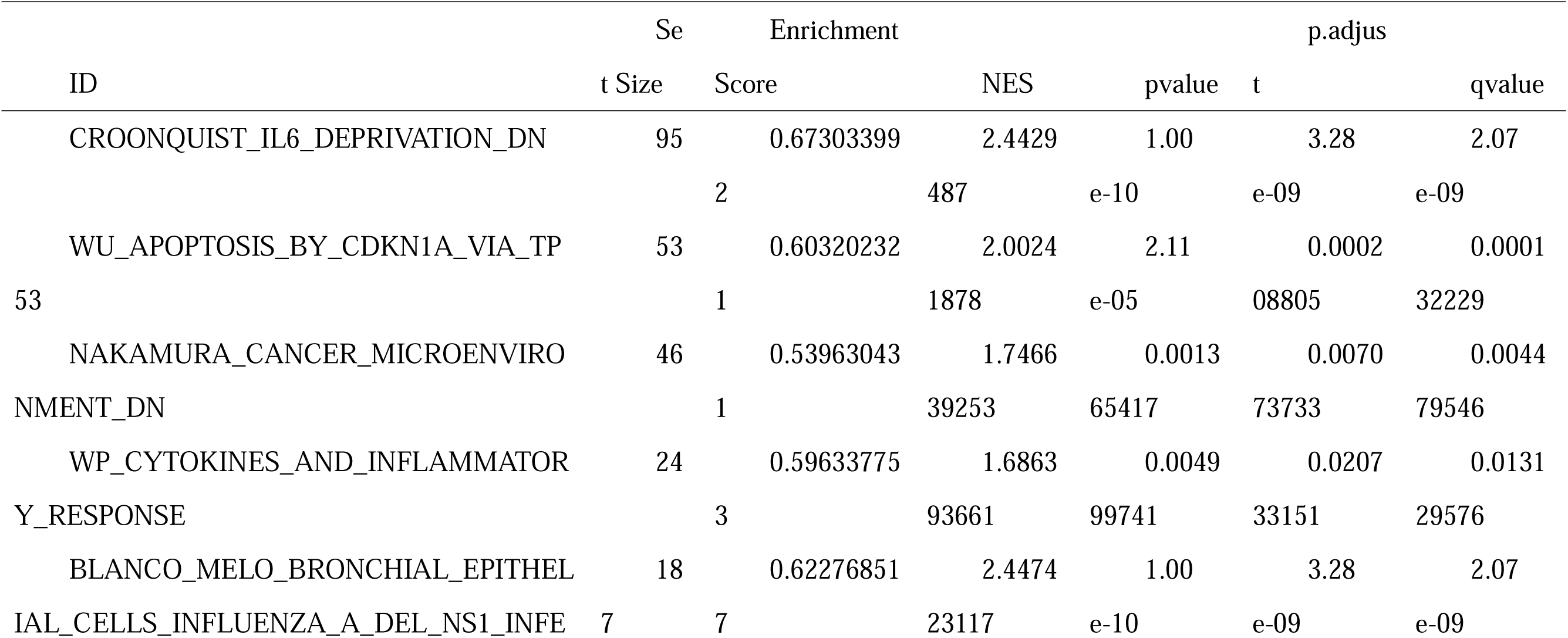
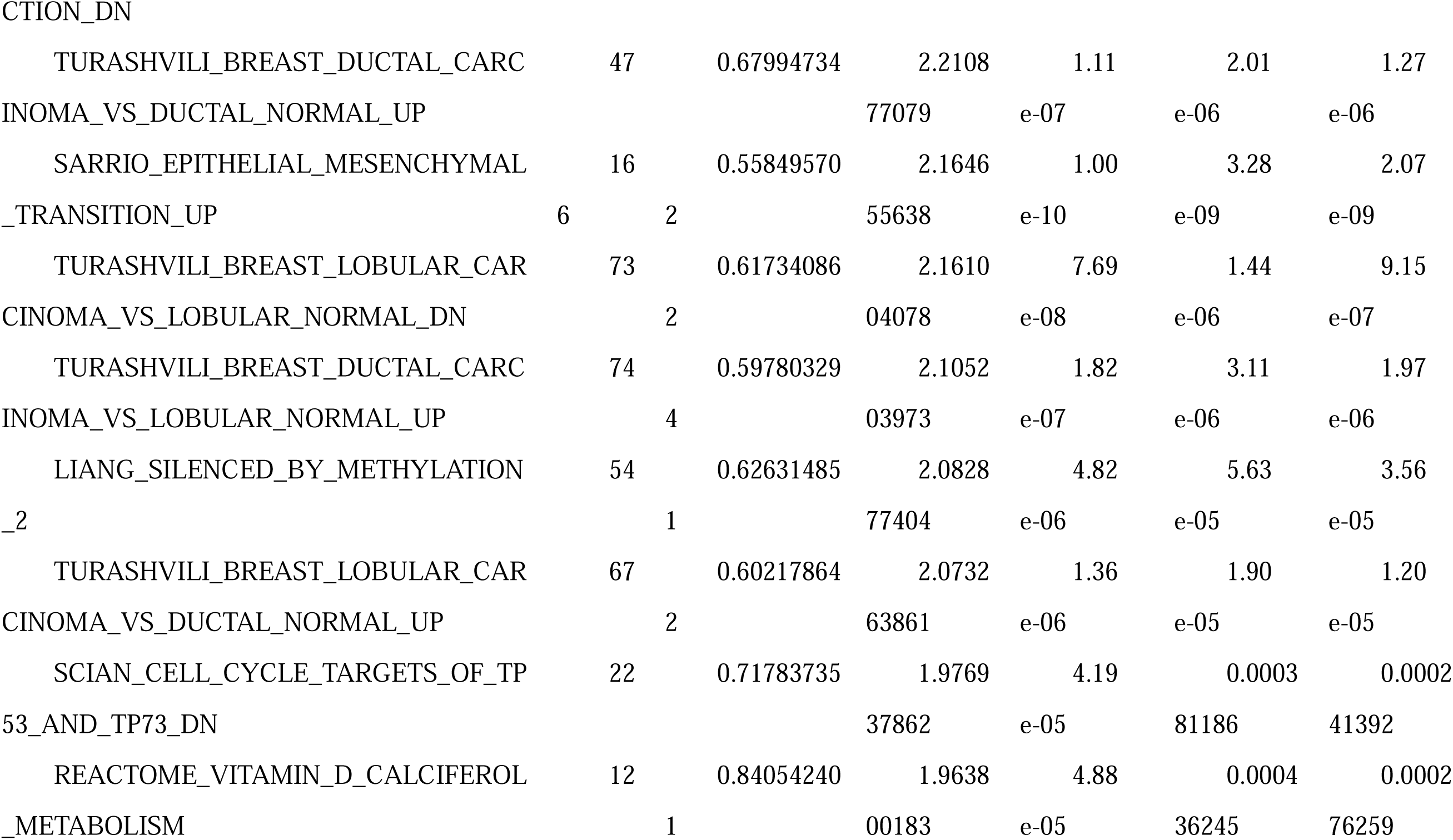
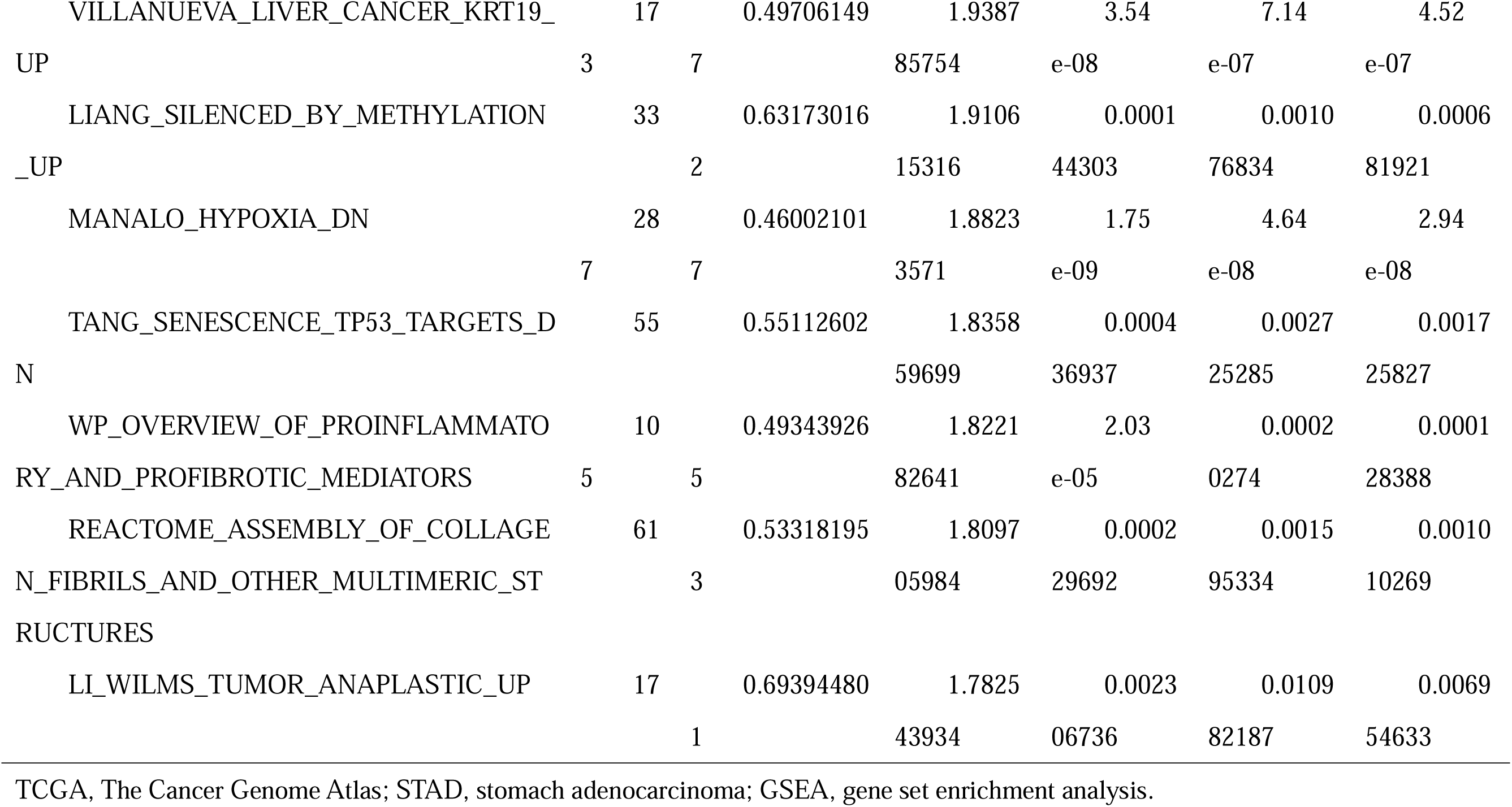
Results of GSEA for TCGA-STAD.

This suggests that inflammatory factors such as IL6 not only promote inflammatory responses, but may also participate in the occurrence and development of GC and the sensitivity regulation of immunotherapy by regulating cell apoptosis, affecting the tumor microenvironment. The abnormal activation of these signaling pathways may contribute to the immune escape and drug resistance of tumor cells, providing a theoretical basis for understanding the molecular mechanisms of GC and optimizing immunotherapy strategies.

### 3.7 Development a gastric cancer prognostic model

To establish a prognostic risk model for STAD, a univariate Cox regression analysis was conducted on 71 GANDIRDEGs. The variables that exhibited a p < 0.10 in the analysis were subsequently illustrated through a forest plot Fig. 7A). The analysis revealed that 12 GANDIRDEGs, including *INHBA, OLR1, ROS1, TIMP1, TUBB6, COPZ2, KCNMB2, EMP3, NMUR1, EPHA5, TACR1*, and *IL6,* were statistically significant different. A LASSO regression analysis was then conducted to evaluate the prognostic value of these DEGs, with the findings illustrated in a LASSO regression model diagram (Fig. 7B) and a LASSO variable trajectory diagram (Fig. 7C). The LASSO regression model ultimately contained six genes: *INHBA, OLR1, ROS1, EPHA5, TACR1,* and *IL6*. A multivariate Cox regression analysis was conducted utilizing these six genes, and a nomogram was created to investigate the associations between the expression level of the LASSO risk score and clinical outcomes (Fig. 7D). The LASSO risk score was calculated according to the subsequent formula:

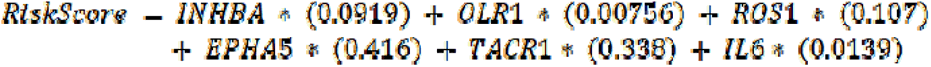

**Fig. 7.**
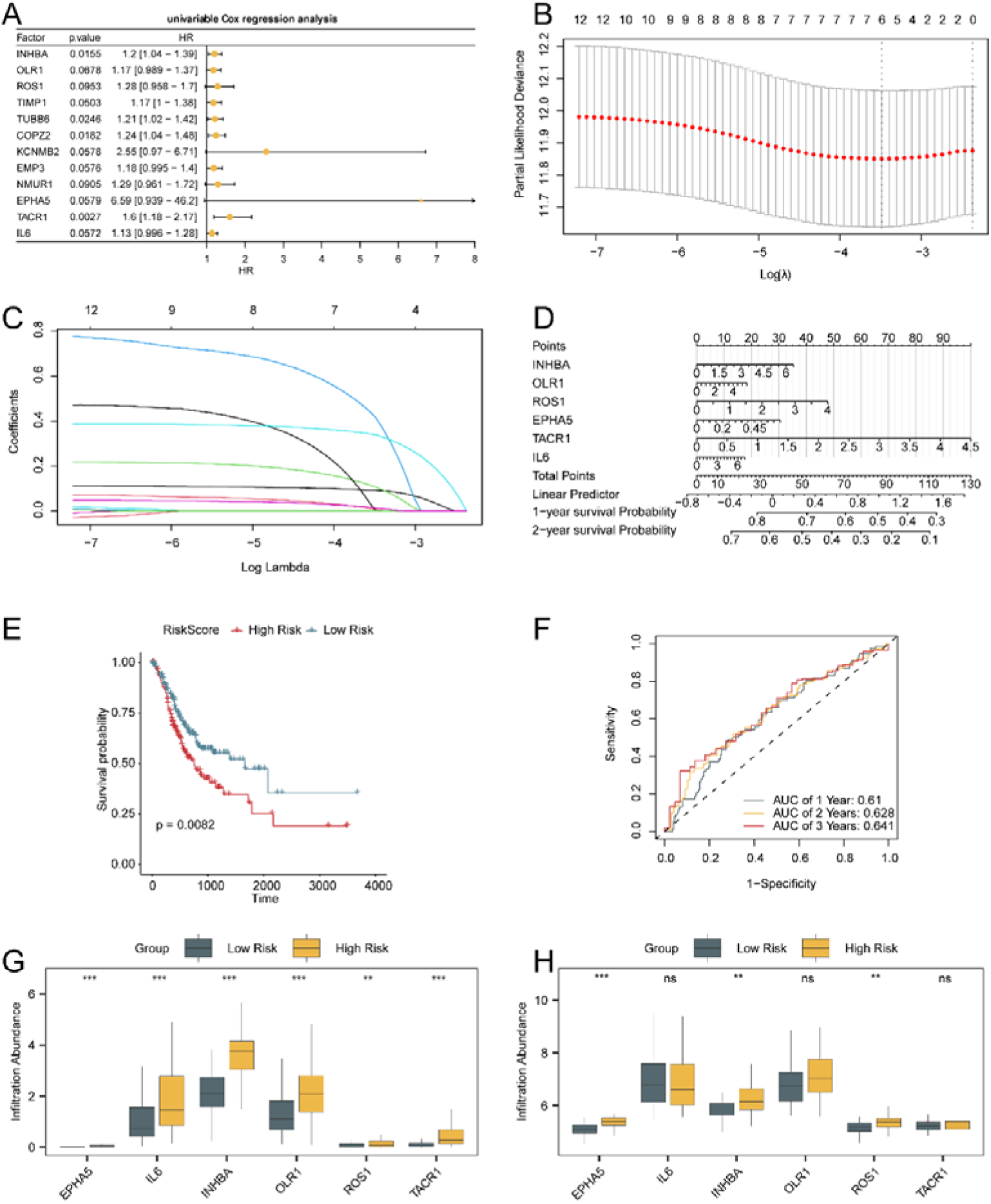
Analysis of LASSO and Cox Regression. **A.** Forest plot of 12 GANDIRDEGs according to the univariate Cox regression model. **B-C.** Plots of the prognostic risk model (B) and variable trajectories (C) from LASSO. **D.** Nomogram of 6 model genes according to the multivariate Cox regression model. **E.** Kaplan-Meier (KM) curves for prognosis between the HR and LR groups according to LASSO and overall survival (OS) in patients with STAD. **F.** Time-dependent ROC curve of the LASSO risk score. **G.** Group comparison plots of the six prognostic risk model-associated genes of TCGA-STAD. **H.** Group comparison plots of the six prognostic risk model-associated genes of STAD samples from the combined GEO datasets. ns stands for p value ≥ 0.05, not statistically significant; ** represents p value < 0.01 and is highly statistically significant; *** represents p value < 0.001, denoting most highly statistically significant; AUC demonstrated limited accuracy, falling between 0.5 and 0.7. The dark green color indicates the LR cohort, whereas the dark yellow color signifies the HR cohort.

To evaluate the diagnostic efficacy of the LASSO risk score in relation to OS, KM curve analysis was conducted utilizing the R package ‘survival’, resulting in KM curves based on the LASSO risk score (Fig. 7E). The analysis revealed a statistically meaningful disparity in OS between the HR and LR cohorts in the TCGA-STAD dataset (p < 0.05). Furthermore, time-dependent ROC curves for 1-, 2-, and 3-year survival were plotted (Fig. 7F), indicating that the LASSO risk score had limited predictive accuracy for prognosis, and the most accurate predictive ability was noted during the third year (AUC= 0.641). GC samples obtained from the TCGA-STAD dataset were classified into HR and LR groups according to the median LASSO risk score. The expression levels of the six model genes were compared between these risk groups (Fig 7G), revealing significant differences in five of the genes (p < 0.001), with all six genes exhibiting elevated expression in the HR cohorts compared with the LR cohorts.

Finally, the risk score for the GC samples from the consolidated GEO datasets was computed using the LASSO risk coefficient, which allowed for the classification of the GC samples into HR and LR cohorts based on the median risk score derived from the LASSO analysis. We drew a comparison chart of the expression levels of six prognostic risk model related genes in the HR and LR cohorts (Fig 7H). The analysis revealed notable statistically significant variations in three of the genes (p < 0.01), whereas five genes presented elevated expression levels in the HR cohort compared with the LR cohort.

### 3.8 Validation of the prognostic model for GC

To validate the accuracy and discrimination of the prognostic risk model for STAD, the LASSO risk score was computed utilizing gene expression levels alongside the LASSO coefficients derived from the TCGA-STAD dataset. Both univariate and multivariate Cox regression analyses were performed to investigate the associations between the LASSO risk score and clinical outcomes. The outcomes of the univariate Cox regression analysis were illustrated via a forest plot (Fig. 8A), which revealed significant correlations between clinical stage, age, risk score, and prognosis. A nomogram was constructed to assess the predictive accuracy of the model (Fig. 8B). The LASSO risk score offered a notably increased utility within the model in comparison with other variables.

**Fig. 8.**
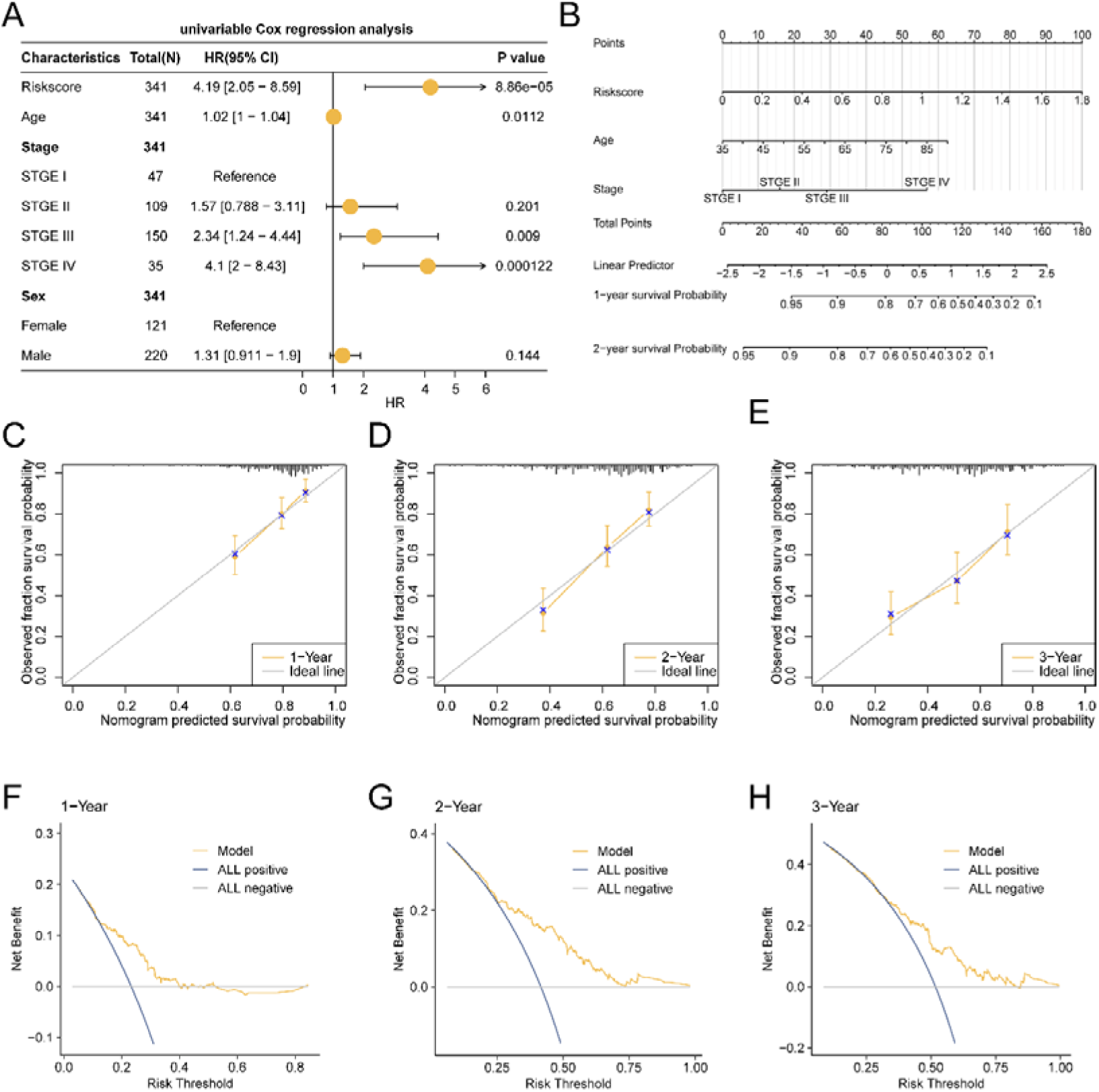
Validation of the prognostic model. **A.** Forest plot of clinical variables: stage, age, sex, and risk score according to the univariate Cox regression model. **B.** Nomograms of stage, age, and the LASSO risk score from the multivariate Cox regression model. **C-E.** Calibration curves of the 1-year (C), 2-year (D), and 3-year (E) prognostic risk models for patients with STAD. **F-H.** 1 -year (F), 2-year (G), and 3-year (H) decision curve analysis (DCA) plots of the prognostic risk model for patients with STAD.

A calibration curve was employed to assess the prediction accuracy of the model for 1-year (Fig. 8C), 2-year (Fig. 8D), and 3-year (Fig. 8E) survival. The calibration curve plots the model-predicted survival probability against the actual survival probability, with closer alignment indicating better prediction accuracy. The findings showed that the STAD prognostic risk model from the TCGA-STAD dataset demonstrated superior clinical predictive accuracy for 1-year survival. Decision curve analysis (DCA) was used to evaluate the clinical applicability of the prognostic risk model for 1-year (Fig. 8F), 2-year (Fig. 8G), and 3-year (Fig. 8H) survival outcomes, revealing that the model’s clinical prediction was most effective for 2-year survival, followed by 3-year and 1-year survival.

### 3.9 Analysis of immune cell infiltration in HR and LR Cohorts

The ssGSEA algorithm was employed to quantify the abundance of infiltrating immune cells among 28 different immune cell types from the TCGA-STAD dataset. A comparative plot was generated to illustrate the variations in the abundance of infiltrating immune cells between the HR and LR cohorts (Fig. 9A). This analysis demonstrated statistically significant differences (p < 0.05) in 22 immune cell types, including activated B cells, activated CD8 T cells, activated dendritic cells, central memory CD4 T cells, central memory CD8 T cells, effector memory CD4 T cells, effector memory CD8 T cells, eosinophils, gamma delta T cells, immature dendritic cells, immature B cells, macrophages, MDSCs, mast cells, memory B cells, natural killer(NK) cells, natural killer T (NKT)cells, plasmacytoid dendritic cells, T follicular helper cells, regulatory T cells, type 1 T helper(Th1) cells, and type 2 T helper(Th2) cells.

**Fig. 9.**
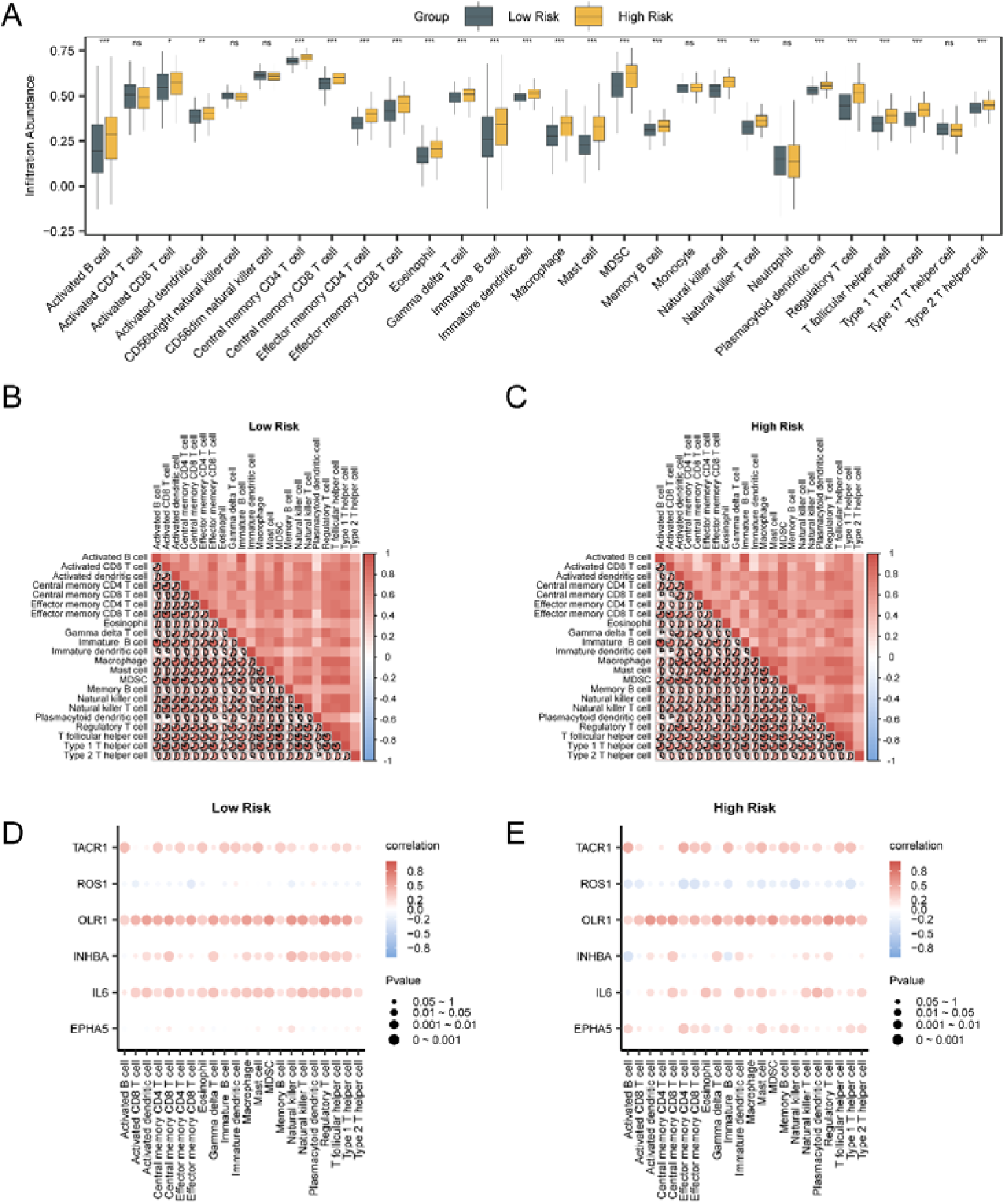
Immune infiltration analysis by ssGSEA algorithm. **A.** Comparison of immune cells in different groups of STAD samples. **B-C.** The findings from the correlation analysis regarding the abundance of infiltrating immune cells in the LR cohort (B) and HR cohort (C) of STAD samples. **D-E.** Bubble plot illustrating the correlation between the abundance of infiltrating immune cell and model-related genes in the LR (D) and HR (E) cohorts of patients with STAD. ns stands for a p value ≥ 0.05, not statistically significant; * represents a p value < 0.05, statistically significant; ** represents a p value < 0.01, highly statistically significant; *** represents a p value < 0.001 and highly statistically significant. The absolute value of the correlation coefficient (r value) below 0.3 was weak or no correlation, that between 0.3 and 0.5 was weak correlation, between 0.5 and 0.8 was moderate correlation, and above 0.8 was strong correlation. The LR group is shown in dark green, and the HR cohort is shown in dark yellow. Red is positively correlated, blue is negatively correlated, and the intensity of the color indicates the magnitude of the correlation.

Correlation analyses of the 22 immune cell infiltration abundances in the GC samples were visualized via a correlation heatmap (Fig 9B-C). The results indicated that the majority of immune cells present in the LR cohort were strongly positively correlated, with the most robust correlation observed between activated B cells and immature B cells (r = 0.941, p < 0.05) (Fig. 9B). Similarly, in the HR cohort, the majority of immune cells also demonstrated strong positive correlations, with the strongest correlation between activated B cells and immature B cells (r = 0.951, p < 0.05) (Fig. 9C). The association between model genes and immune cell infiltration abundance was illustrated through a correlation bubble plot (Fig. 9D-E). The findings indicated that the majority of immune cells in the two groups had strong positive correlations, with the strongest association between the *OLR1* gene and regulatory T cells (r= 0.652, p < 0.05) (Fig. 9E).

### 3.10 TIDE, MSI, TMB, immune checkpoint, and immunophenoscore (IPS) analysis

The samples obtained from the dataset were classified into HR and LR categories on the basis of the median risk score derived from LASSO analysis. The TIDE algorithm was employed to evaluate the sensitivity of GC samples to immunotherapy, and the results were depicted through a comparative group plot. (Fig. 10A). The results indicated a statistically significant difference in TIDE immunotherapy scores between the HR and LR cohorts (p < 0.001), with the LR cohort exhibiting lower scores than the HR cohort did, suggesting that the LR cohorts may have a more favorable response to immunotherapy.

**Fig.10.**
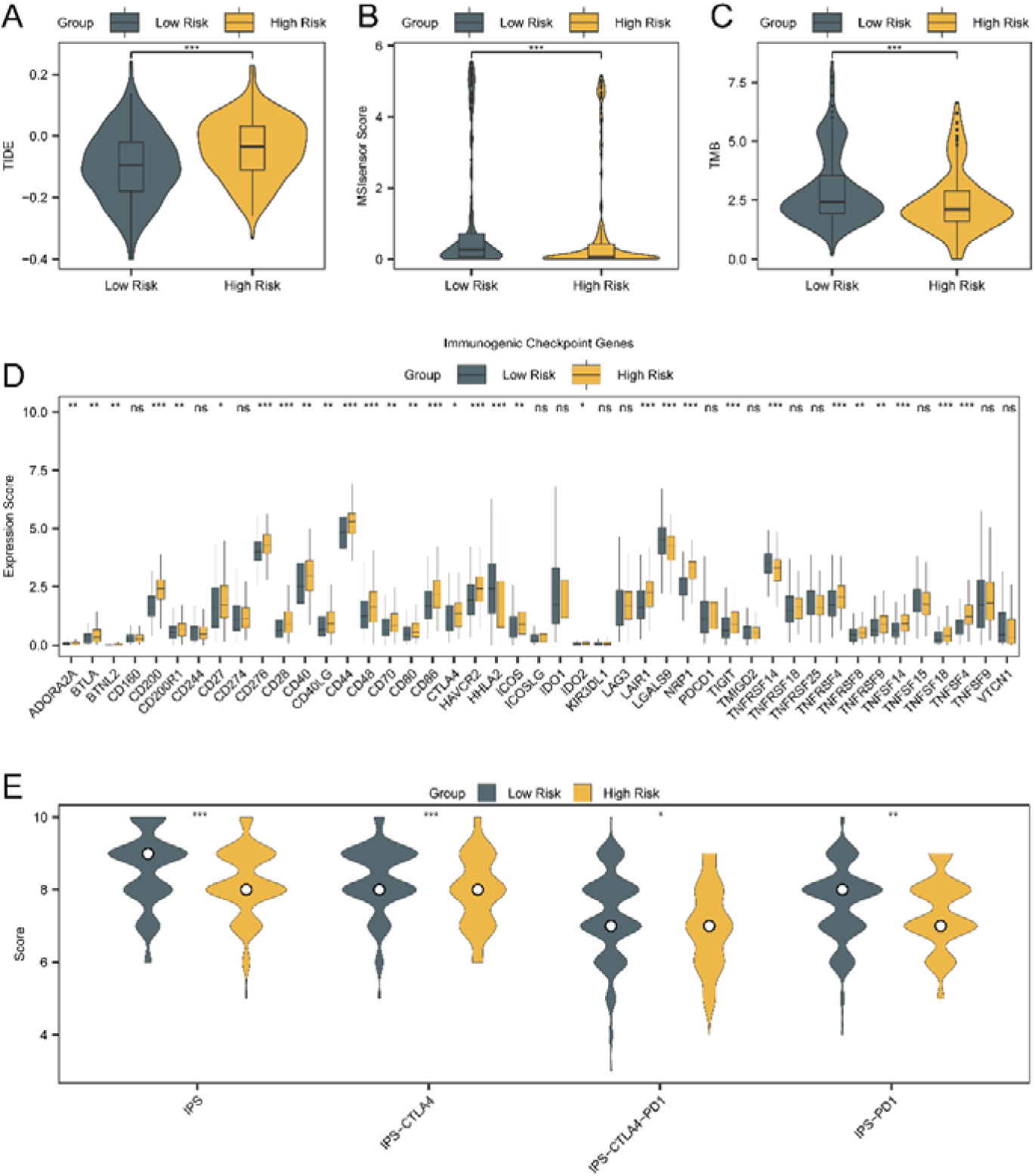
TIDE, MSI, TMB, immune checkpoint genes and IPS analysis. **A-D.** TIDE (A), MSI (B), TMB (C), and immune checkpoint genes (D) group comparison graphs between HR and LR cohorts of STAD samples from the TCGA-STAD dataset. **E.** Group comparison plots of different immunogenicity score (IPS) groups in the HR group and LR group of patients with STAD samples. * signifies a p-value of less than 0.05, indicating statistical significance; ** indicates a p-value of less than 0.01, which is considered highly statistically significant; and *** denotes a p-value of less than 0.001, reflecting the highest level of statistical significance. Dark green represents the LR cohorts and dark yellow represents the HR cohorts.

The differences in microsatellite instability (MSI) and tumor mutational burden (TMB) scores between different cohorts were analyzed (Fig. 10B-C). The findings revealed statistically meaningful differences in the MSI (p < 0.001) and TMB (p < 0.001) scores between the two groups, with the LR group having higher scores than the HR group did.

To obtain immune checkpoint genes (ICGs) from the literature, we conducted an intersection analysis between the identified ICGs and the genes available in the dataset. We generated a matrix composed of 45 ICGs and their expression levels, with comprehensive details provided in Table S4. The variations in the expression levels of 45 ICGs between the two cohorts were analyzed utilizing the Mann-Whitney U test (Fig. 10D). The results revealed statistically significant differences (p < 0.05) for 30 ICGs between the two cohorts (Table S5).

The immunophenoscore (IPS) for the GC samples was obtained from the TCIA database, and group comparisons of the IPS between the HR and LR cohorts were visualized via the R package ggplot2 (Fig. 10E). The results showed statistically significant differences in IPS, IPS-PD1, IPS-CTLA4, and IPS-CTLA4-PD1 between the two groups (p < 0.01).

### 3.11 Drug sensitivity analysis

Drug sensitivity analysis was conducted via the Genomics of Drug Sensitivity in Cancer (GDSC) database, the Cancer Cell Line Encyclopedia (CCLE) database, and the CellMiner database. Utilizing the pRRophetic algorithm, the expression levels of specific model genes, along with data on drug activity, were employed to predict the sensitivity of these model genes to commonly utilized anticancer drugs. The results were visualized via correlation plots (Fig. 11A-C). The findings revealed that 55 interacting drugs were identified in the GDSC database (Fig. 11A), with *EPHA5* showing the strongest correlation with all drugs. In the CCLE database, 15 interacting drugs were identified (Fig11. B), with *EPHA5* showing the strongest correlation with all drugs. In the CellMiner database, 17 interacting drugs were identified (Fig. 11C), with most model genes showing positive correlations with all drugs.

**Fig.11.**
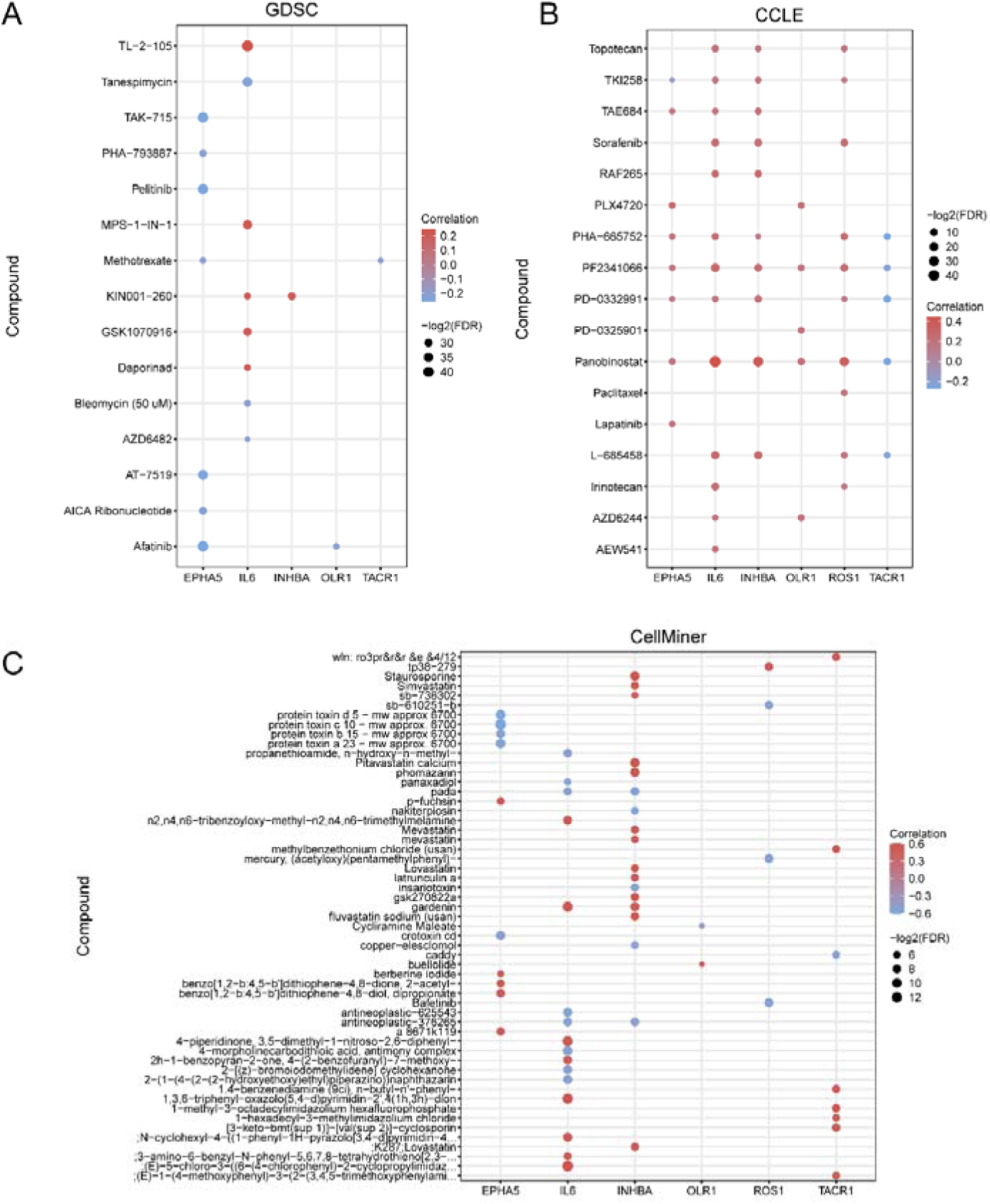
Drug sensitivity analysis. Drug sensitivity analysis results of model genes from the GDSC database **(A)**, CCLE database **(B)** and CellMiner database **(C)** are presented. An absolute correlation coefficient (r value) of less than 0.3 suggests a weak or negligible relationship, whereas values ranging from 0.3 to 0.5 indicate a weak correlation. Furthermore, coefficients between 0.5 and 0.8 reflect a moderate correlation, and those exceeding 0.8 represent a strong correlation. In this context, red is used to denote a positive correlation, whereas blue is employed to represent a negative correlation

## 4 Discussion

GC is a widely occurring malignant tumor and is the fifth most common cause of cancer-associated death globally. In China, GC morbidity and mortality rank second, as stated by the National Cancer Institute of China [40]. The prognosis for patients diagnosed with GC remains unfavorable, characterized by a limited five-year survival rate. Most GC cases are identified at more advanced stages, resulting in restricted therapeutic alternatives and an disease progression. Current therapeutic methods for advanced GC, such as chemotherapy and chemoembolization, are suboptimal, underscoring the pressing necessity for novel diagnostic and therapeutic targets. This study revealed significant alterations in gene expression associated with glycosylation and inflammatory responses in GC, which may be closely associated with the disease’s biological behavior. By integrating data from multiple databases, we established a prognostic risk model for GC, on the basis of six pivotal genes highlighted through LASSO regression analysis. This model not only predicts overall survival in GC patients but also demonstrates good predictive accuracy through time-dependent ROC curve analysis. The AUC of the model in overall survival prediction is about 0.64, indicating moderate predictive ability. This result suggests that the model has certain risk stratification and survival prediction value, but has not yet reached the high accuracy level required for clinical application.

The identified six-gene signature includes *INHBA, OLR1, ROS1, EPHA5, TACR1, and IL6.* Inhibin subunit beta A (INHBA), a constituent of the transforming growth factor-beta (TGF-β) superfamily, is critically involved in several cancer types [41]. Both INHBA and its homodimer activin A exert multifaceted effects on the regulation of immune responses and tumor progression [42]. Elevated INHBA expression in GC patients has been associated with worse outcomes [43]. Liu et al. demonstrated that FAP^+^ GC mesenchymal stromal cells enhanced GC progression by releasing the cytokine INHBA and remodeling the extracellular matrix (ECM), providing a theoretical basis for targeted therapies aimed at the tumor stroma in GC [44]. OLR1, considered a pancancer prognostic marker, is marked overexpression in cancer-associated fibroblasts (CAFs) and is closely linked with adverse prognoses in lung cancer and several other malignancies [45]. Wu et al. reported that patients exhibiting high levels of OLR1 expression are more susceptible to immune evasion and derive fewer benefits from immune checkpoint inhibitor (ICI) therapies [46]. Nevertheless, the specific role of OLR1 in GC remains understudied. In our investigation, OLR1 expression was associated with OS in patients with GC, but the specific mechanisms involved require further investigation. The ROS1 proto-oncogene encodes a receptor tyrosine kinase that is structurally homologous to other oncogenic drivers and has been recognized as a potential therapeutic target in lung cancer. Research has showed that approximately 35% of patients with *ROS1*-positive non-small cell lung cancer (NSCLC) develop metastases to the central nervous system (CNS) [47,48]. Erythropoietin-producing human hepatocellular (Eph) receptors constitute the most extensive category of receptor tyrosine kinases (RTKs). Research indicates that EphA5 is differentially expressed in GC and is significantly associated with the Lauren classification, lymph node metastasis, HER2 expression, and Ki-67 expression [49]. Tachykinin receptor 1 (TACR1) encodes the receptor for substance P, also known as neurokinin-1. This protein plays a mediating role in the phosphatidylinositol metabolism of substance P. Multiple studies have indicated that the activation of TACR1 (by binding with substance P) is linked to tumor angiogenesis and cellular proliferation across various cancer types [50,51]. Interleukin-6 (IL6) encodes a cytokine involved in inflammation and B-cell maturation. It also acts as an endogenous pyrogen, inducing fever in patients with autoimmune diseases or infections. Liu et al. demonstrated that lower baseline levels of serum IL-6 correlate with prolonged progression-free survival (PFS) in NSCLC patients [52]. The IL-6/JAK/STAT3 signaling pathway is pivotal in GC progression [53].

By utilizing these six genes, patients were stratified into HR and LR cohorts, with significant differences in OS observed between the groups. A nomogram was developed through multivariate regression analysis, showing that the LASSO risk score exhibited markedly greater effectiveness within the model than the other variables did. Notably, most genes in our risk model are related to immune responses, aligning with the current focus on immunotherapy in GC. Furthermore, some genes in the model exhibited statistically significant discrepancies in expression levels across different groups, suggesting their potential key roles in GC prognosis. Immune cell infiltration analysis further unveiled variations in the immune microenvironment between the HR and LR cohorts, thereby offering fresh perspectives on the mechanisms of immune evasion pertinent to GC. As a core component of the immunosuppressive microenvironment, the enrichment of Tregs in HR groups may promote immune escape of tumor cells, inhibit the function of effector T cells, and weaken the body’s immune surveillance of tumors. MDSCs also show higher infiltration in HR groups, and these cells further enhance tumor immune tolerance and are closely related to immune therapy resistance by secreting immunosuppressive factors, inhibiting T cell activation, and other mechanisms. On the contrary, CD8+T cells, as the main anti-tumor effector cells, have significantly higher infiltration levels in the LR group, indicating that patients in the LR group have stronger anti-tumor immune response ability, which may be related to better prognosis and sensitivity to immunotherapy. These findings not only reveal differences in the immune microenvironment between HR and LR groups, but also provide a theoretical basis for understanding the mechanisms of immune escape and immune therapy response in gastric cancer.

Our findings from the drug sensitivity analysis, indicate that the expression levels of genes featured in our model correlate with sensitivity to various anticancer agents, suggesting potential biomarkers for the pharmacological treatment of GC. Additionally, our study indicates that disparities in the expression levels of genes associated with immune checkpoints across different cohorts may influence patient responses to immune checkpoint blockade therapies [54]. These results provide a theoretical basis for the subsequent development of chemotherapy or targeted therapy strategies based on molecular subtyping, and also suggest that some model genes can serve as molecular markers for predicting drug sensitivity. In practical clinical practice, targeting the expression levels of key genes such as EPHA5, IL6, and ROS1 may help optimize drug selection and improve treatment benefits for GC patients.

Traditional single pathway analysis often fails to fully reflect the complex biological characteristics of tumors. However, this study, by integrating glycosylation and inflammation related genes, can more systematically reveal the dynamic changes in the tumor microenvironment and its impact on immune escape and drug sensitivity. Joint analysis not only enhances the biological explanatory power of the model in patient prognosis stratification and immune microenvironment feature identification, but also provides more clinically valuable molecular basis for personalized treatment. The integrated model constructed in this study demonstrates unique advantages in revealing the immune infiltration status and predicting drug sensitivity in gastric cancer patients, laying the foundation for precision medicine and the development of new treatment strategies.

Although our study offers valuable insights, it also presents certain limitations. Primarily, the analysis relies on information obtained from publicly accessible databases, and the heterogeneity of data sources may lead to certain biases. Although different datasets have been integrated through standardization and batch effect removal methods, the potential impact of batch effects on the analysis results cannot be completely ruled out. Additionally, the risk model has not been independently validated in a prospective clinical cohort and lacks functional validation at the laboratory level (such as immunohistochemistry, quantitative PCR, etc.). Therefore, the clinical applicability and molecular mechanisms of the model still need to be further confirmed in real-world sample and experimental studies. Furthermore, future studies should investigate the molecular mechanisms underlying these genes in GC development and their impact on treatment response.

## 5 Conclusion

In conclusion, this study successfully identified six molecular markers (*INHBA, OLR1, ROS1, EPHA5, TACR1,* and *IL6*) that are associated with the prognosis of GC via a thorough examination of gene expression data. We have also developed a prognostic risk assessment model that provides significant insights for the precision treatment of GC and lays a foundation for future clinical research and drug development endeavors.

## Acknowledgements

The authors would like to acknowledge the TCGA database (https://tcga-data.nci.nih.gov/tcga) and GEO databases (https://www.ncbi.nlm.nih.gov/geo/) for providing their platforms and those contributors for uploading their valuable datasets.

## Author contributions

Kun Yang, Zhaopu Li and Jiaqing Lin designed the study; Zhaopu Li, Jiaqing Lin, Weipeng Liu, Yang Li and Wei Zhu collected and analyzed the data; Kun Yang and Shixiong Yang drafted the manuscript. All authors read and approved the final version.

## Data availability

The data and materials used to support this study’s findings are available from the corresponding author upon request or public databases

## Declarations

### Ethics approval and consent to participate

This study was approved by the Ethics Committee of Xiaogan Hospital Affiliated to Wuhan University of Science and Technology. The research was conducted in accordance with the International Conference and the Declaration of Helsinki.

### Consent to Publish

All authors have consented to the publication of this manuscript.

### Clinical trial

Not applicable

